# Drug-adapted cancer cell lines reveal drug-induced heterogeneity and enable the identification of biomarker candidates for the acquired resistance setting

**DOI:** 10.1101/2020.02.13.947374

**Authors:** Martin Michaelis, Mark N. Wass, Ian Reddin, Yvonne Voges, Florian Rothweiler, Stephanie Hehlgans, Jaroslav Cinatl, Marco Mernberger, Andrea Nist, Thorsten Stiewe, Franz Rödel, Jindrich Cinatl

## Abstract

Survivin is a drug target and the survivin suppressant YM155 a drug candidate for high-risk neuroblastoma. Findings from one YM155-adapted subline of the neuroblastoma cell line UKF-NB-3 had suggested that increased ABCB1 (mediates YM155 efflux) levels, decreased SLC35F2 (mediates YM155 uptake) levels, decreased survivin levels, and *TP53* mutations indicate YM155 resistance. Here, the investigation of ten additional YM155-adapted UKF-NB-3 sublines only confirmed the roles of ABCB1 and SLC35F2. However, cellular ABCB1 and SLC35F2 levels did not indicate YM155 sensitivity in YM155-naïve cells, as indicated by drug response data derived from the Cancer Therapeutics Response Portal (CTRP) and the Genomics of Drug Sensitivity in Cancer (GDSC) databases. Moreover, the resistant sublines were characterised by a remarkable heterogeneity. Only seven sublines developed on-target resistance as indicated by resistance to RNAi-mediated survivin depletion. The sublines also varied in their response to other anti-cancer drugs. In conclusion, cancer cell populations of limited intrinsic heterogeneity can develop various resistance phenotypes in response to treatment. Therefore, individualised therapies will require monitoring of cancer cell evolution in response to treatment. Moreover, biomarkers can indicate resistance formation in the acquired resistance setting, even when they are not predictive in the intrinsic resistance setting.

## Introduction

YM155 (sepantronium bromide) was introduced as an anti-cancer drug candidate that inhibits expression of the *BIRC5* gene, which encodes the protein survivin [1]. In the meantime, YM155 has been suggested to exert additional and/ or alternative mechanisms of anti-cancer actions including induction of DNA damage, inhibition of NFκB signaling, induction of death receptor 5 expression and/ or suppression of MCL-1, XIAP, cIAP-1/2, BCL-2, BCL-XL, FLIP, and/ or EGFR [2-12].

A number of studies have investigated the potential of YM155 against neuroblastoma cells [13-16]. Neuroblastoma is the most common extracranial solid childhood tumor. Treatment outcomes in high-risk neuroblastoma patients remain unsatisfactory. About 50% of these patients relapse and have a 5-year-survial rate below 10% [17-20]. We have recently shown that suppression of survivin expression is the main mechanism through which YM155 exerts its anti-neuroblastoma effects [15]. Notably, the New Drug Development Strategy (NDDS, a project of Innovative Therapies for Children with Cancer, the European Network for Cancer Research in Children and Adolescents, and the International Society of Paediatric Oncology Europe Neuroblastoma) has categorized survivin as a high priority drug target in neuroblastoma and YM155 as a high priority drug [21].

The formation of acquired resistance is a central problem in (metastasised) cancer diseases that need to be treated by systemic drug therapy. Although many cancers initially respond well to therapy, resistance formation is common and cures are rare [22]. Hence, biomarkers that indicate early therapy failure are needed to adapt therapies if resistance emerges. Liquid biopsies (e.g. circulating tumor cells) enable the monitoring of cancer cell evolution in patients with ever more detail [23]. However, the translation of the resulting information into improved therapies is hampered by a lack of understanding of the processes underlying acquired resistance formation and, in turn, a lack of biomarkers.

Most studies focus on biomarkers that indicate whether a certain cancer cell (population) is likely to respond to a certain treatment, but not on biomarkers that indicate early that a current therapy has stopped working. This also applies to the previous studies that investigated the efficacy of YM155 in neuroblastoma [13,14,16]. However, it is known that intrinsic and acquired resistance mechanisms may substantially differ [24-26]. Using a single YM155-adapted neuroblastoma cell line, we identified increased ABCB1 (also known as P-glycoprotein or MDR1) expression, decreased SLC35F2 (Solute Carrier Family 35 Member F2) expression, decreased survivin expression, and loss-of-p53-function as potential markers of resistance formation to YM155 [15]. Given the tremendous (intra-tumour) heterogeneity in cancer [27], it is likely that the processes, which result in acquired resistance formation, are equally complex. If so, then a larger number of models of acquired resistance to a certain drug will be needed to adequately address the complexity of the resistance formation process.

To test this hypothesis, we here established and characterised 10 further YM155-adapted UKF-NB-3 neuroblastoma cell lines. To see whether we can obtain information from our acquired resistance models that cannot be identified from traditional approaches using non-adapted cell lines, we also analysed YM155 response data from the two large pharmacogenomics screens Genomics of Drug Sensitivity in Cancer (GDSC) and Cancer Therapeutic Response Portal (CTRP) [28,29]. We found a remarkable heterogeneity between the individual sublines, although they all had been derived from the same parental cell line. An increase in cellular ABCB1 levels and/ or a decrease in SLC35F2 levels indicate resistance formation to YM155, although the ABCB1 and/ or SLC35F2 levels cannot be used to infer YM155 sensitivity in YM155-naïve cell lines. The use of the panel of YM155-adapted cell lines further enabled us to show that the cellular survivin levels and the *TP53* status do not reliably indicate resistance formation.

## Results

### YM155-adapted UKF-NB-3 sublines display pronounced YM155 resistance

Representative photos of the morphology of the project cell lines are presented in Figure S1 and the doubling times in Table S1. All YM155-adapted UKF-NB-3 sublines displayed pronounced YM155 resistance (Figure 1, Table S1). The relative resistance expressed as fold change of the YM155 IC50 values in the YM155-adapted UKF-NB-3 sublines divided by the YM155 IC50 value in UKF-NB-3 (IC50: 0.55nM) ranged between 38 (UKF-NB-3^r^YM155^20nM^IV; IC50: 21.0nM) and 76 (UKF-NB-3^r^YM155^20nM^VI; IC50: 41.9nM) (Table S1). The fold changes of the YM155 IC90 values in the YM155-adapted UKF-NB-3 sublines relative to the YM155 IC90 value in UKF-NB-3 (IC90: 1.01nM) ranged from 30 (UKF-NB-3^r^YM155^20nM^IV; IC90: 29.8nM) to 135 (UKF-NB-3^r^YM155^20nM^VI; IC90: 136nM) (Table S1).

**Figure 1.**
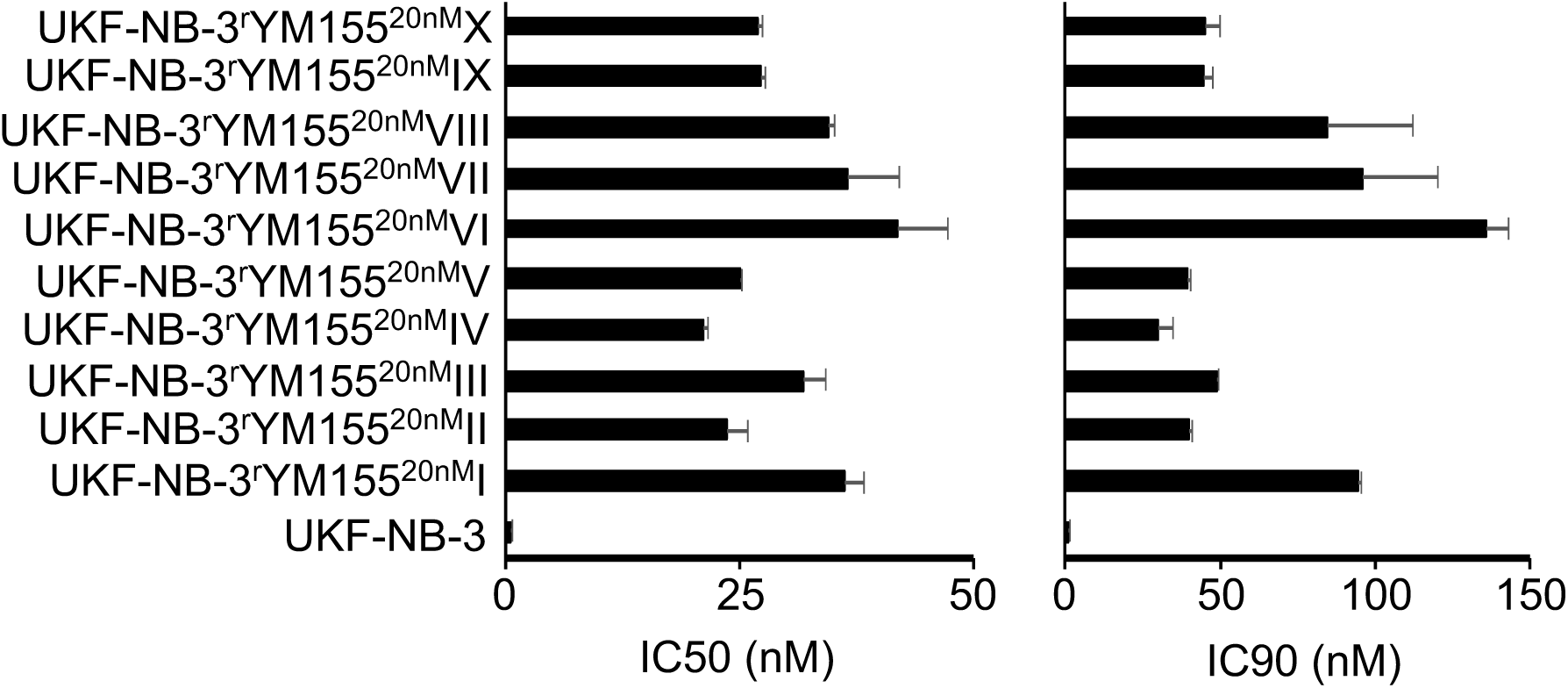
YM155 concentrations that reduce the viability of UKF-NB-3 and its YM155-adapted sublines by 50% (IC50) or 90% (IC90) as determined by MTT assay after 120h of incubation. Numerical values are presented in Table S1. * P < 0.05 relative to UKF-NB-3

### The cellular *TP53* status is not a reliable indicator of YM155 sensitivity

Originally, the cellular *TP53* status was described to not directly influence the anti-cancer action of YM155 [30]. In agreement, the analysis of the Genomics of Drug Sensitivity in Cancer (GDSC) and Cancer Therapeutics Response Portal (CTRP) databases did not indicate differences in YM155 sensitivity between cell lines in dependence on their *TP53* status (wild-type or mutant) (Figure 2).

**Figure 2.**
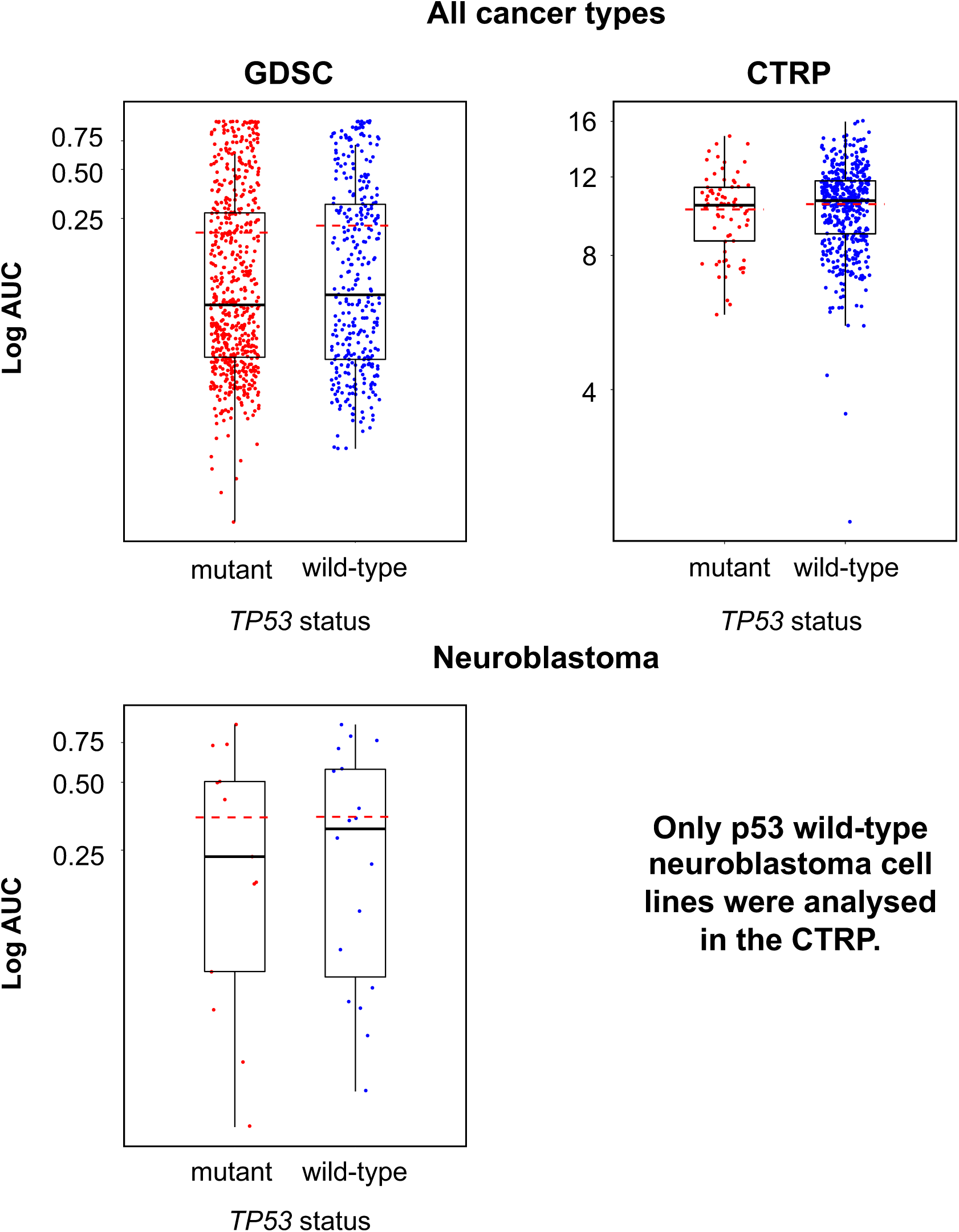
YM155 sensitivity in p53 wild-type and p53 mutant cancer cell lines based on the analysis of GDSC and CTRP data, both determined across all investigated cancer types/ cell lines (GDSC, p = 0.458; CTRP, p = 0.216) and in a neuroblastoma-specific analysis (GDSC, p = 0.922; all 12 neuroblastoma cell lines in the CTRP harbour wild-type *TP53*).

However, the activation of p53 signalling seems to be involved in the anticancer mechanism of action of YM155 at least in some cancer cells. We have previously shown in neuroblastoma cells that YM155 activates p53 signalling, that p53 activation using MDM2 inhibitors enhances the YM155 effects, and that p53 depletion reduces cancer cell sensitivity to YM155 [15]. In addition, a YM155-adapted UKF-NB-3 subline harboured a *TP53* mutation [15]. However, all 10 YM155-adapted UKF-NB-3 sublines that we investigated here displayed wild-type *TP53* as indicated by *TP53* next generation sequencing. The cellular p53 levels did also not differ consistently between UKF-NB-3 and its YM155-resistant sublines (Figure S2). The YM155-resistant UKF-NB-3 sublines remained similarly sensitive to the MDM2 inhibitor and p53 activator nutlin-3 as UKF-NB-3 (Table S2). Hence, our findings do not suggest that YM155 adaption is generally associated with loss of p53 function in neuroblastoma cells. The cellular *TP53* status is not a reliable indicator of YM155 sensitivity, neither in the intrinsic nor in the acquired resistance setting.

### Cellular survivin levels do not reliably indicate YM155 response

Some studies suggested cancer cells with high survivin levels to be particularly sensitive to YM155 [30-32]. However, other studies failed to detect an association between the cellular survivin status and YM155 activity [15,33]. When we compared the YM155 sensitivity between cancer cell lines with high and low survivin expression, we found statistically significant differences across all cell lines in the GDSC and CTRP datasets, but not the neuroblastoma cell lines (Figure 3). It was not possible to predict whether a certain cell line was sensitive to YM155 based on the cellular survivin level (Figure 3).

**Figure 3.**
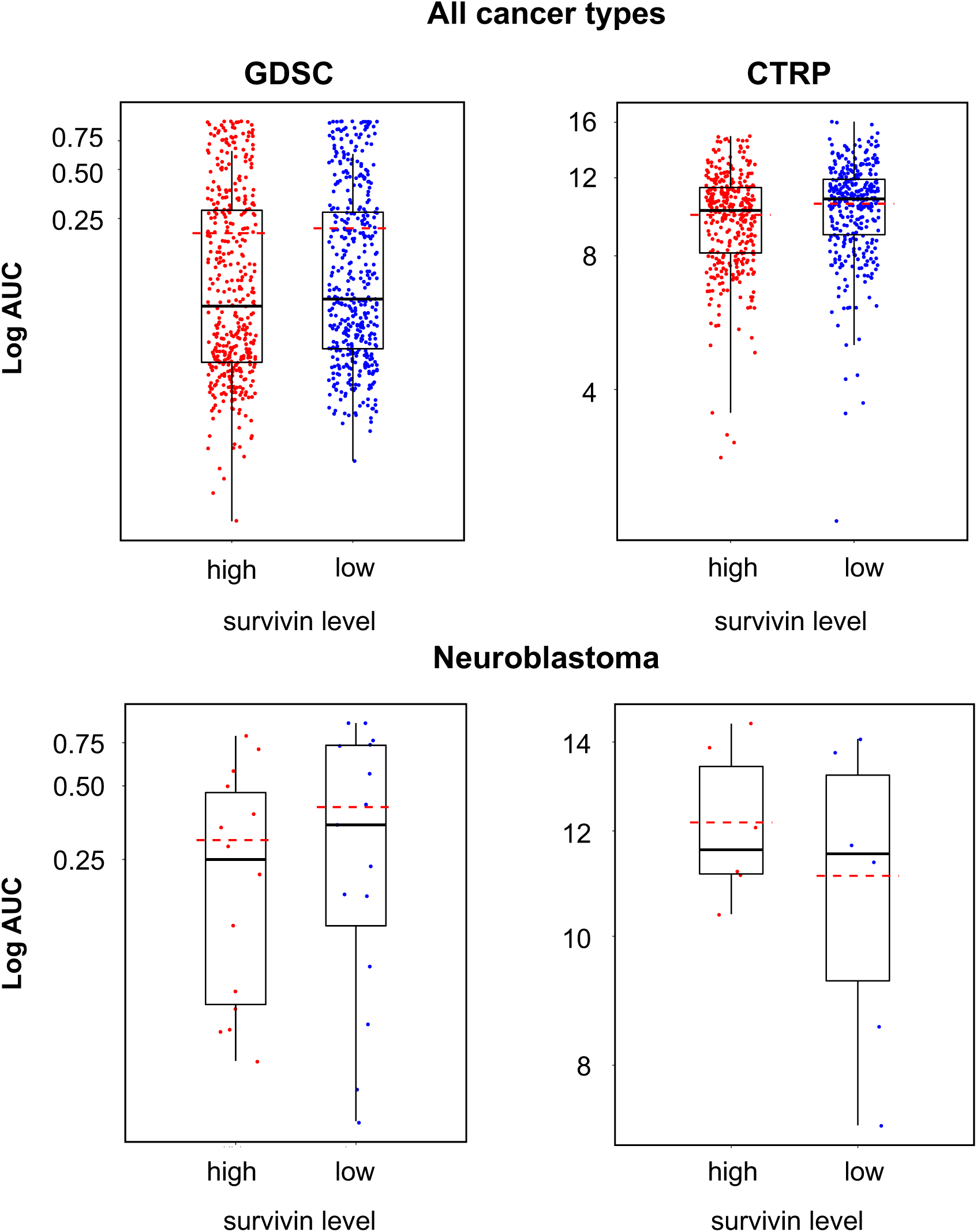
YM155 sensitivity in cell lines characterised by high or low survivin expression based on the analysis of GDSC and CTRP data, both determined across all investigated cancer types/ cell lines (GDSC, p = 0.048; CTRP, p < 0.001) and in a neuroblastoma-specific analysis (GDSC, p = 0.425; CTRP, p = 0.699).

Notably, a YM155-adapted UKF-NB-3 subline had previously displayed reduced survivin levels relative to the parental cell line [15]. However, the analysis of the 10 additional YM155-adapted UKF-NB-3 sublines in this study revealed that resistance acquisition to YM155 was not associated with a consistent change in the survivin expression patterns (Figure S3).

### Acquired YM155 resistance is associated with decreased sensitivity to survivin depletion

Our previous findings had suggested that YM155 predominantly exerts its anti-neuroblastoma effects via suppression of survivin expression [15]. Seven of the ten YM155-adapted UKF-NB-3 sublines (I, III, V, VII, VIII, IX, X) displayed decreased sensitivity to siRNA-mediated survivin depletion. Two sublines were similar sensitive as parental UKF-NB-3 cells (II, VI), and one subline (IV) was more sensitive (Figure 4, Figure S4). This shows that a majority of the YM155-resistant cell lines have developed on-target resistance. It also indicates that the YM155 resistance mechanisms differ between the individual UKF-NB-3 sublines.

**Figure 4.**
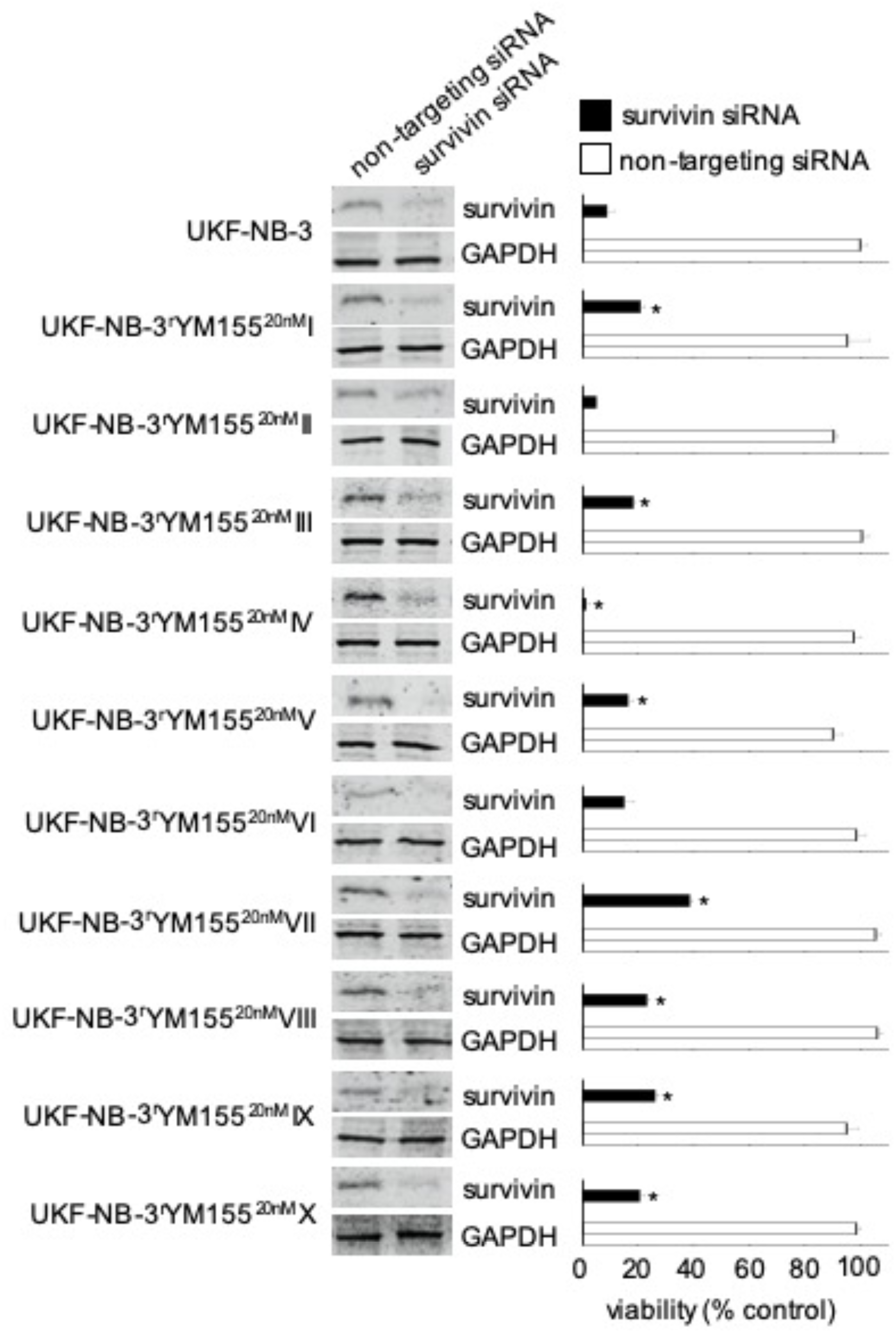
Effects of siRNA-mediated BIRC5/ survivin depletion on the viability of UKF-NB-3 and its YM155-adapted sublines. Western blots confirm reduced survivin levels 48h post transfection. Viability of cells transduced with siRNA directed against BIRC5/ survivin or non-targeting siRNA was determined relative to untreated control cells 168h post transfection by MTT assay. * P < 0.05 relative to untreated cells

### Relevance of cellular ABCB1 and SLC35F2 levels in the context of YM155 resistance

Increased cellular ABCB1 (mediates YM155 efflux) levels and decreased SLC35F2 (mediates cellular YM155 uptake) levels have previously been identified as important YM155 resistance mechanisms [13,15,16,34]. To further investigate the relationship between ABCB1 and SLC35F2 levels and YM155 sensitivity, we compared the YM155 sensitivity in cell lines that displayed low or high expression of the respective genes using GDSC and CTRP data. In agreement with previous data, high *ABCB1* expression (Figure 5) and low *SLC35F2* expression (Figure 6) were associated with reduced YM155 sensitivity. When we used transcriptomics data from the GDSC and CTRP to correlate the expression of all genes with YM155 sensitivity, *ABCB1* ranked as the gene whose expression was most strongly correlated to the YM155 AUC (area under the curve, unit used to quantify drug response) (Table 1) in the GDSC and CTRP. *SLC35F2* expression was most strongly inversely correlated to the YM155 AUC (Table 2) in both data sets. There were no further overlaps among the top 10 genes between the two databases (Table 1, Table 2). However, the YM155 sensitivity of a certain cell line could not be reliably predicted based on the cellular ABCB1 and/ or SLC35F2 levels (Figure 5, Figure 6).

**Table 1.**
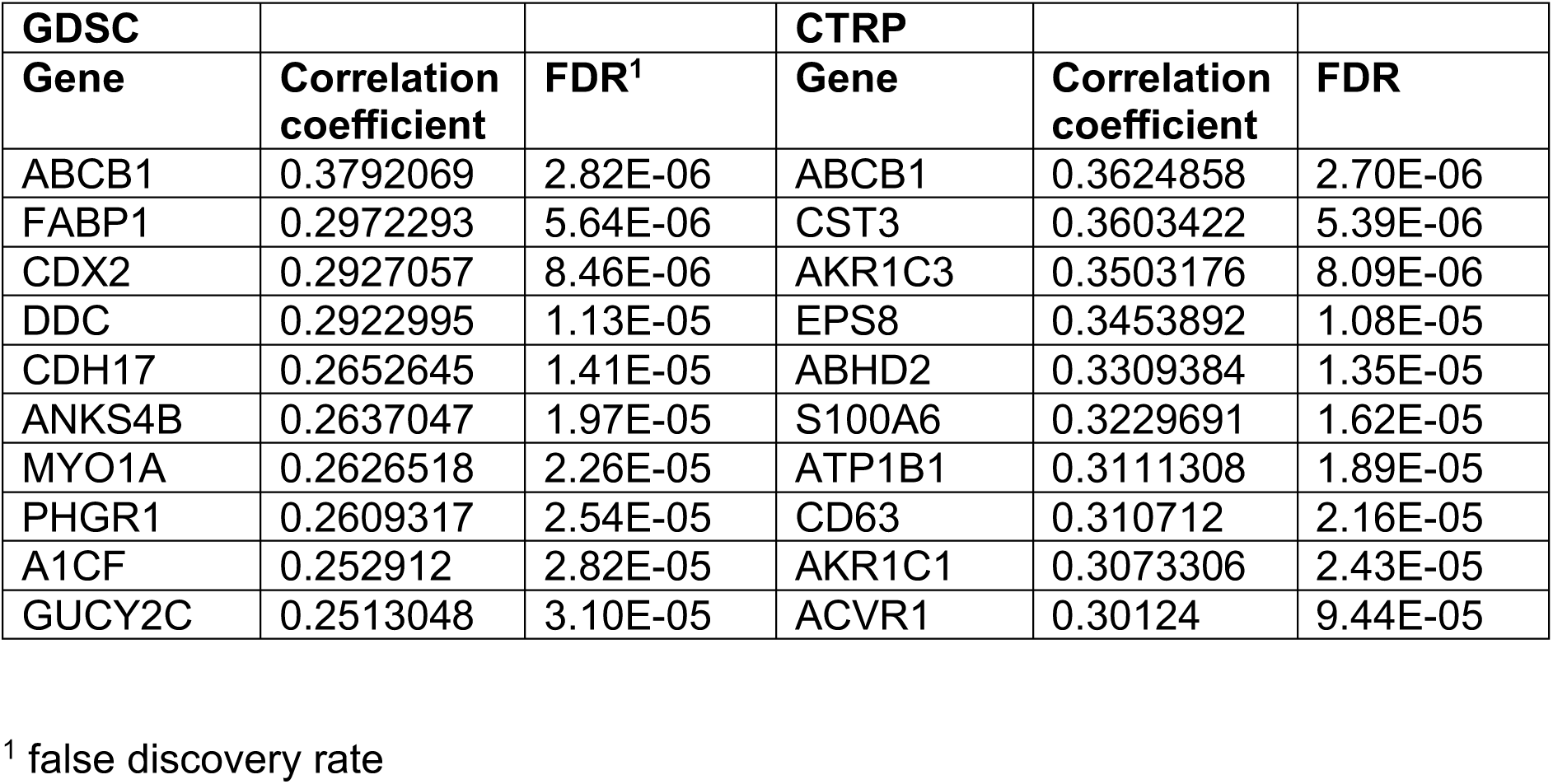
Top 10 genes whose expression is most strongly correlated with the YM155 AUC in the Genomics of Drug Sensitivity in Cancer (GDSC) database and the Cancer Therapeutics Response Portal (CTRP) as indicated by the Pearson Correlation Coefficient.

**Table 2.** Top 10 genes whose expression is most strongly inversely correlated with the YM155 AUC in the Genomics of Drug Sensitivity in Cancer (GDSC) database and the Cancer Therapeutics Response Portal (CTRP) as indicated by the Pearson Correlation Coefficient.

| **GDSC** |  |  | **CTRP** |  |  |
| --- | --- | --- | --- | --- | --- |
| **Gene** | **Correlation coefficient** | **FDR<sup>1</sup>** | **Gene** | **Correlation coefficient** | **FDR** |
| SLC35F2 | -0.2643809 | 1.69E-05 | SLC35F2 | -0.3868041 | 9.17E-05 |
| ALKBH8 | -0.2042557 | 0.000121 | CD19 | -0.349156 | 8.90E-05 |
| CWF19L2 | -0.1926243 | 0.000180 | CD79B | -0.346035 | 8.63E-05 |
| RCSD1 | -0.1855956 | 0.000226 | SPIB | -0.341887 | 8.36E-05 |
| P2RY8 | -0.1834937 | 0.000257 | SNX22 | -0.338893 | 8.09E-05 |
| RGS19 | -0.1831031 | 0.000259 | TCL1A | -0.338393 | 7.82E-05 |
| FLI1 | -0.1825892 | 0.000268 | LOC100130458 | -0.337940 | 7.55E-05 |
| VAV1 | -0.1819569 | 0.000273 | BLK | -0.330471 | 7.28E-05 |
| ATM | -0.1816322 | 0.000276 | CD79A | -0.324465 | 7.01E-05 |
| ARHGAP19 | -0.1813872 | 0.000282 | VPREB3 | -0.322878 | 6.74E-05 |

<sup>1</sup> false discovery rate

**Figure 5.**
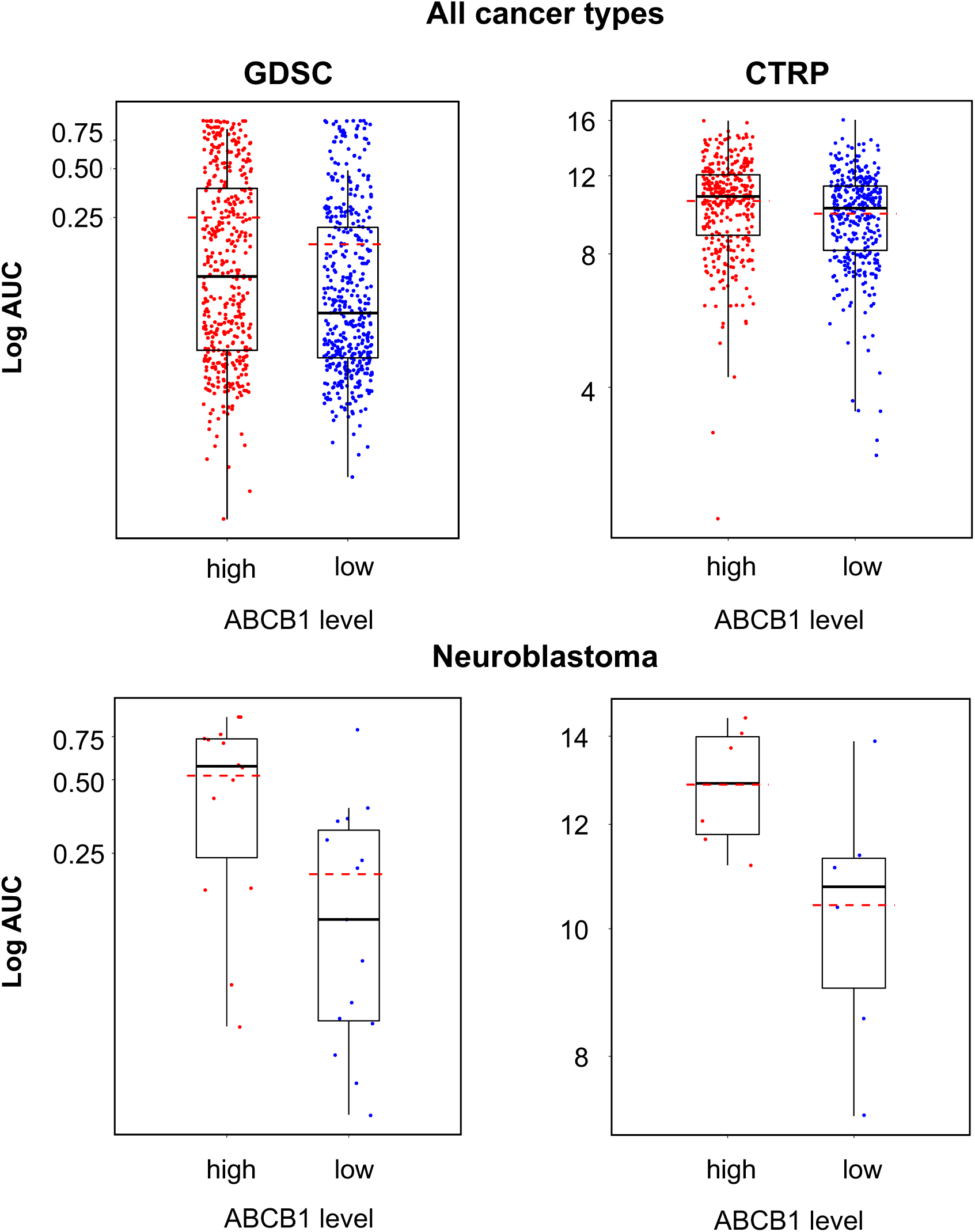
YM155 sensitivity in cancer cell lines characterised by high or low ABCB1 expression based on the analysis of GDSC and CTRP data, both determined across all investigated cancer types/ cell lines (GDSC, p < 0.001; CTRP, p < 0.001) and in a neuroblastoma-specific analysis (GDSC, p = 0.006; CTRP, p = 0.04).

**Figure 6.**
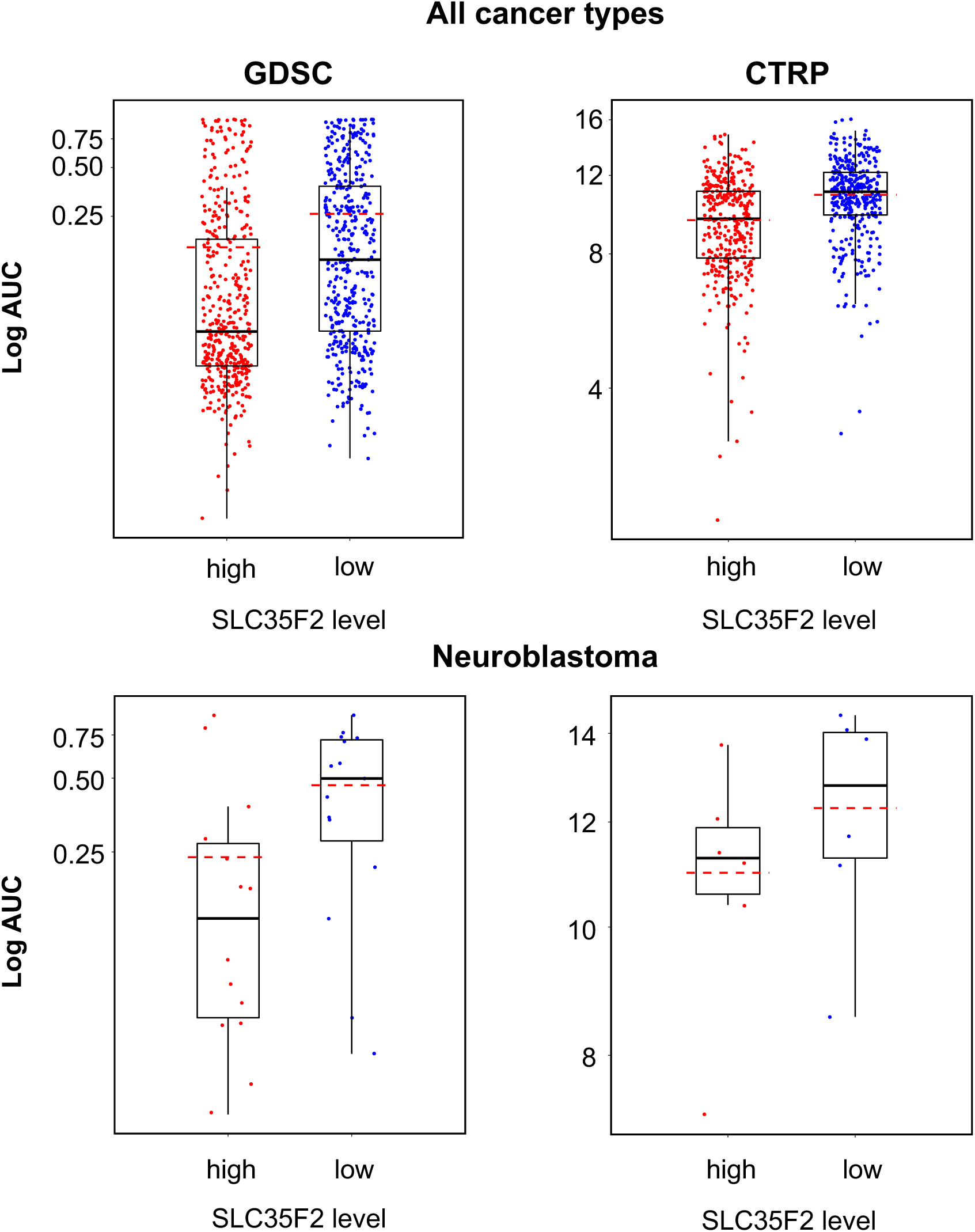
YM155 sensitivity in cancer cell lines characterised by high or low SLC35F2 expression based on the analysis of GDSC and CTRP data, both determined across all investigated cancer types/ cell lines (GDSC, p < 0.001; CTRP, p < 0.001) and in a neuroblastoma-specific analysis (GDSC, p = 0.033; CTRP, p = 0.310).

All YM155-adapted UKF-NB-3 sublines displayed increased ABCB1 levels relative to UKF-NB-3 (Figure 7, Figure S5). Acquired YM155 resistance was also generally associated with decreased SLC35F2 levels, in particular in the sublines I, IV, VI, and X (Figure 7, Figure S5). This indicates that increased ABCB1 levels and decreased SLC35F2 levels have potential as biomarkers indicating YM155 resistance formation in response to YM155-based therapies, although cellular ABCB1 and SLC35F2 levels do not enable the prediction of YM155 sensitivity in YM155-naïve cells.

**Figure 7.**
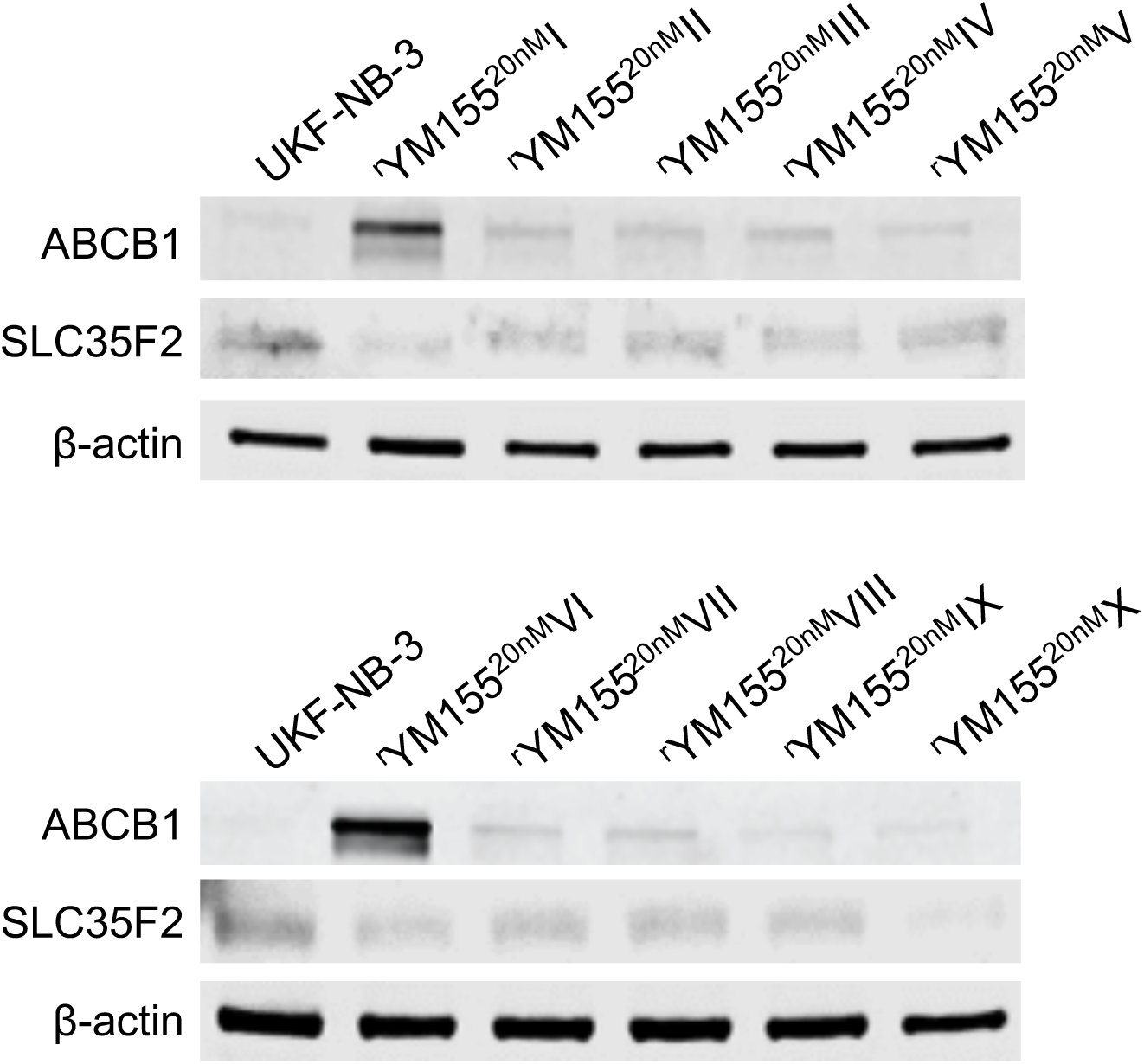
Representative Western blots indicating cellular levels of ABCB1 and SLC35F2 in UKF-NB-3 and YM155-adapted UKF-NB-3 sublines.

### YM155-adapted UKF-NB-3 cells remain sensitive to DNA damage caused by irradiation and cytotoxic drugs

YM155 has been proposed to exert its anti-cancer effects via the induction of DNA damage in some experimental systems [3,5,34,35]. To study whether the acquisition of YM155 resistance was associated with a generally increased resistance to DNA damage, UKF-NB-3 and its YM155-adapted UKF-NB-3 sublines were irradiated at a dose range of one to five Gy. None of the YM155-adapted UKF-NB-3 sublines displayed substantially reduced sensitivity to irradiation relative to UKF-NB-3 (Figure 8). Moreover, none of the YM155-resistant UKF-NB-3 sublines displayed reduced sensitivity to cisplatin (causes DNA crosslinks) or topotecan (topoisomerase I inhibitor), which cause DNA damage by different mechanisms (Figure 8, Table S2). There was also no coherent increase in resistance to the nucleoside analogue gemcitabine (Figure 8, Table S2). These data do not suggest a dominant role of DNA damage induction in the course of the anti-neuroblastoma activity of YM155 in UKF-NB-3 cells.

**Figure 8.**
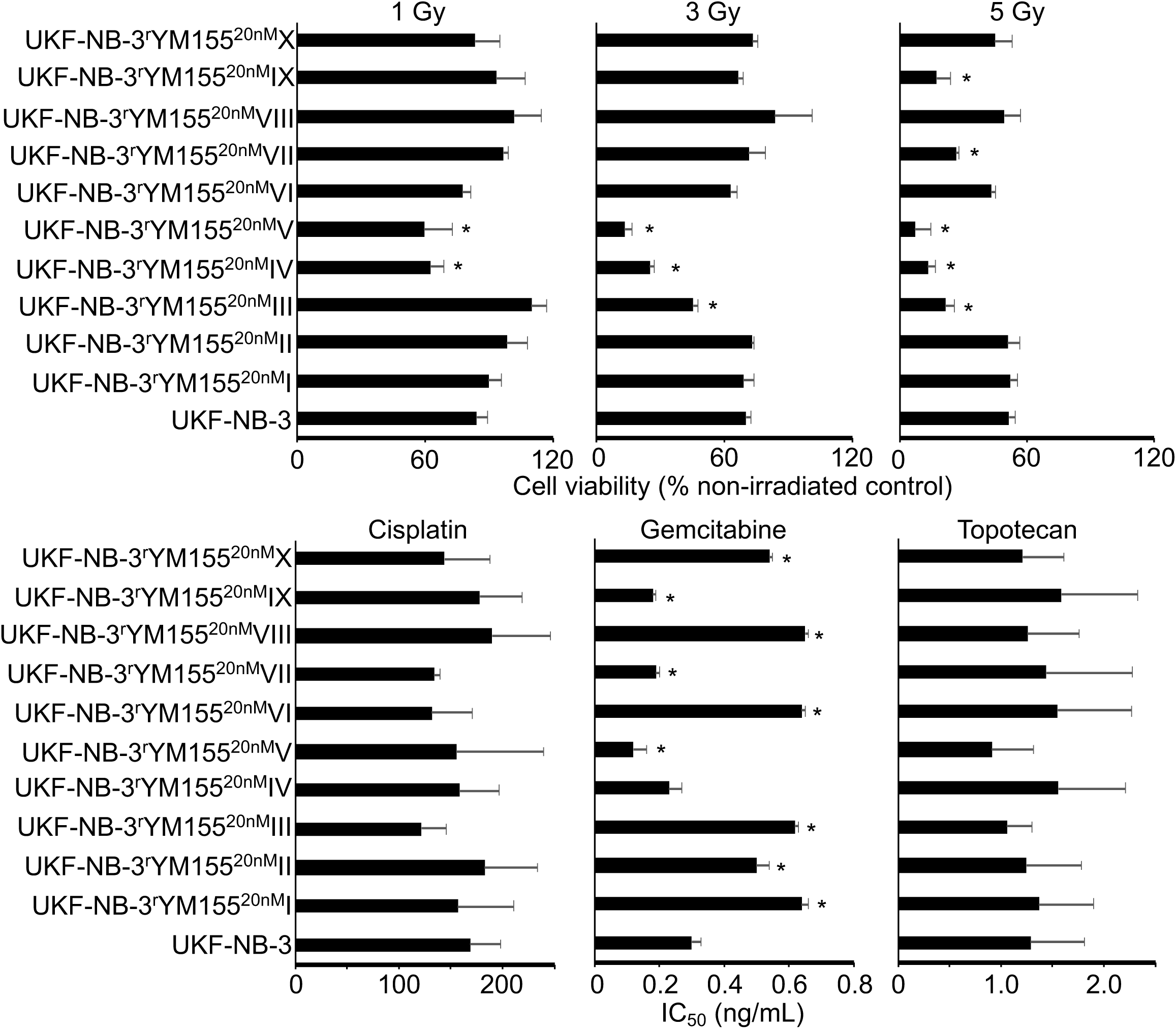
Sensitivity of UKF-NB-3 and its YM155-adapted sublines to irradiation and DNA damaging drugs. The radiation response was determined 72h after irradiation with 1, 3, or 5Gy by MTT assay. Drug concentrations that reduce cell viability by 50% (IC_50_) were determined by MTT assay after 120h of incubation. * P < 0.05 relative to UKF-NB-3

### Heterogeneity among YM155-adapted UKF-NB-3 sublines

While the YM155-adapted UKF-NB-3 sublines displayed limited heterogeneity in response to treatment with cisplatin and topotecan, remarkable differences in the gemcitabine IC_50_s were detected (Figure 8, Table S2). The fold difference between the YM155-adapted subline with the lowest gemcitabine IC_50_ (V, 0.12ng/mL) and the subline with the highest IC_50_ (VIII, 0.65ng/mL) was 5.4-fold. This heterogeneity is in agreement with the up to 29-fold difference observed in cell viability in response to BIRC5/ survivin depletion between the most sensitive (II) and the most resistant (VII) subline (Figure 4). Resistance profiles to the destabilising tubulin-binding agent vincristine also revealed a substantial heterogeneity between the YM155-resistant UKF-NB-3 sublines (Figure 9, Table S2), resulting in a fold difference of 127 between subline VI (vincristine IC_50_: 714ng/mL) and subline IX (vincristine IC_50_: 5.6ng/mL) (Table S2).

**Figure 9.**
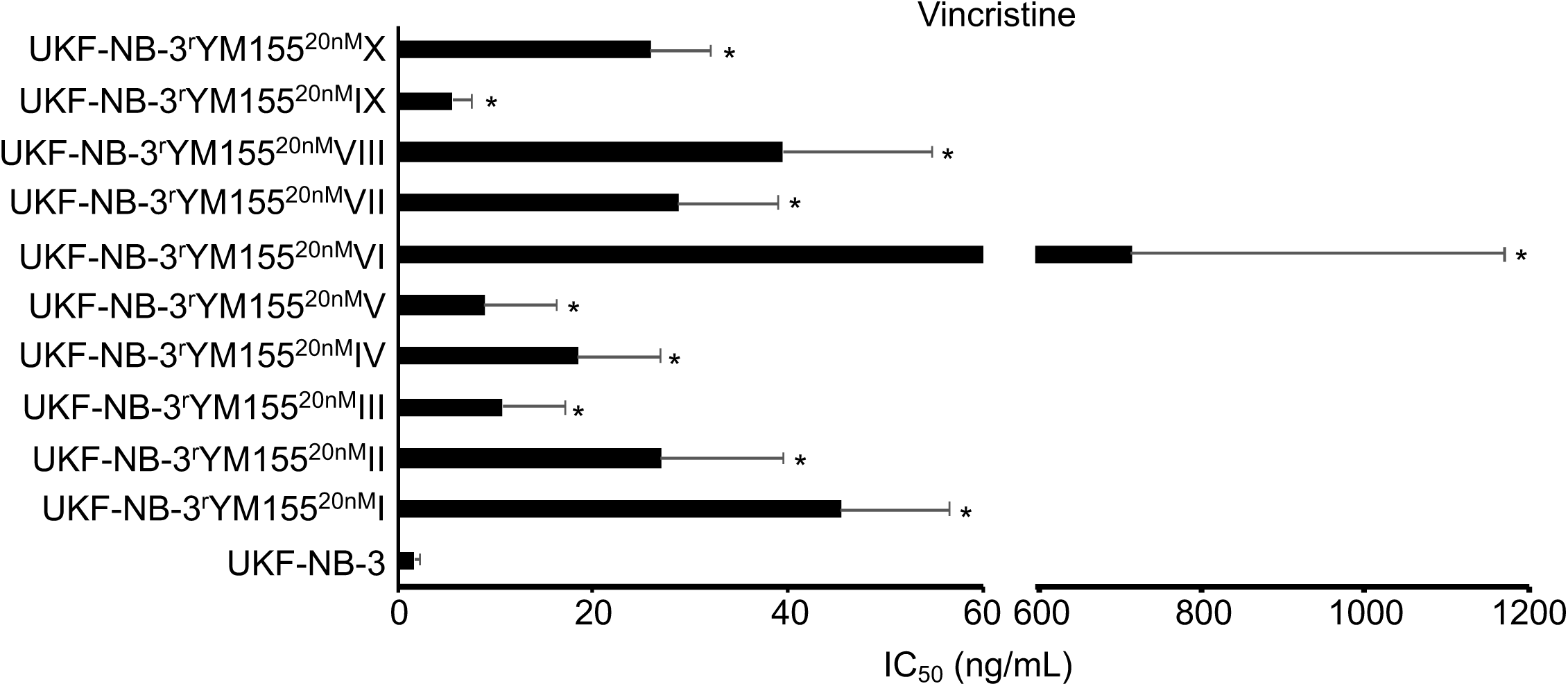
Vincristine concentrations that reduce cell viability by 50% (IC_50_) were determined by MTT assay after 120h of incubation. * P < 0.05 relative to UKF-NB-3

## Discussion

In a previous study, a YM155-adapted subline of the neuroblastoma cell line UKF-NB-3 was characterised by increased cellular ABCB1 levels, decreased SLC35F2 and survivin levels, and a *TP53* mutation [15]. Here, we systematically investigated the relevance of cellular ABCB1, SLC35F2, and survivin levels as well as the *TP53* status as potential biomarkers of YM155 resistance formation in the intrinsic resistance setting, using data derived from the GDSC and CTRP databases [28,29], and in the acquired resistance setting, using an additional set of ten YM155-adapted UKF-NB-3 sublines, which were established in independent experiments.

Increased ABCB1 expression (mediates YM155 efflux) and decreased SLC35F2 expression (mediates cellular YM155 uptake) were identified as YM155 resistance mechanisms in panels of YM155-naïve cell lines that displayed varying levels of these proteins and in functional studies [13,15,16,34], which was further supported by our analysis of GDSC and CTRP data [28,29]. Despite their roles in determining YM155 resistance, however, cellular ABCB1 or SLC35F2 levels did not enable the prediction of whether an individual cell line would be sensitive to YM155 or not. The YM155-adapted UKF-NB-3 cell lines generally displayed elevated cellular ABCB1 levels and reduced SLC35F2 levels relative to UKF-NB-3. Hence, an increase in the cellular ABCB1 levels and/ or a decrease in the SLC35F2 levels have potential as biomarkers that indicate resistance formation, even though the respective cellular levels do not reliably predict YM155 response in YM155-naïve cells.

Initially, the *TP53* status was reported not to influence the anti-cancer effects of YM155 [30], which was further supported by our analysis of GDSC and CTRP data. In neuroblastoma cells, however, YM155 induced p53 signalling, p53 depletion reduced YM155 sensitivity, and a YM155-adapted UKF-NB-3 subline harboured a *TP53* mutation [15]. Here, all ten YM155-adapted UKF-NB-3 sublines retained wild-type *TP53*. Thus, the role of p53 seems to depend on the individual cellular context. Neither the cellular *TP53* status nor the formation of *TP53* mutations can currently be considered as valid biomarkers for YM155 therapies.

The relevance of cellular survivin levels for cancer cell sensitivity to YM155 is not clear [15,30-33]. Our analysis of GDSC and CTRP data indicated that high survivin (BIRC5) expression was associated with increased YM155 sensitivity. However, it was not possible to infer the YM155 sensitivity of a particular cell line based on its survivin status. Reasons for this may include that survivin is not in all cell lines the major therapeutic target of YM155 as it is in neuroblastoma cells [1-12,15,35] and/ or that off-target resistance mechanisms such as ABCB1 and SLC35F2 expression may affect YM155 efficacy independently of the survivin status [13,15,16,34].

The YM155-adapted UKF-NB-3 sublines displayed various survivin levels, demonstrating that resistance formation to YM155 is also not associated with a consistent change in cellular survivin levels. Seven of the YM155-adapted cell lines displayed on-target resistance as indicated by reduced sensitivity to RNAi-mediated BIRC5/ survivin depletion relative to parental UKF-NB-3 cells, further confirming that survivin is a target of YM155 in neuroblastoma cells. However, cellular survivin levels do not represent a reliable biomarker of resistance formation to YM155.

While YM155 was described to act via the induction of DNA damage in some cancer types [3,5,34,35], our previous results did not indicate a causative role of DNA damage induction in the anti-cancer effects of YM155 against neuroblastoma cells [15]. YM155 resistance formation in the YM155-adapted neuroblastoma cell lines was also not associated with generally decreased sensitivity to radiation or DNA damage caused by cisplatin (causes DNA crosslinks), gemcitabine (nucleoside analogue), or topotecan (topoisomerase I inhibitor). This indicates that YM155 resistance formation in neuroblastoma cells is not generally associated with an increased resistance to DNA damage induction.

In this study, the use of multiple models of acquired resistance enabled insights that could not be gained from just one drug-adapted subline. The previous investigation of one YM155-resistant UKF-NB-3 subline had suggested that changes in the cellular *TP53* status and survivin levels indicate resistance formation [15], which was not confirmed in our current panel of ten YM155-adapted UKF-NB-3 sublines. Moreover, the use of multiple sublines provided a pioneering glimpse onto the remarkable heterogeneity of the resistance formation process, even though all resistant sublines were derived from the same parental cell line. Only seven of the ten sublines developed on-target resistance mechanisms as indicated by reduced sensitivity to survivin depletion. The sublines also showed substantial variation in their sensitivity to irradiation (up to 7-fold difference at 5Gy), gemcitabine (up to 5-fold), and vincristine (up to 127-fold). Notably, a much higher heterogeneity would be expected in the clinical situation, in which tumours are already characterised by much higher heterogeneity than cancer cell lines and in which combination therapies are common.

In conclusion, our data reveal a high phenotypic heterogeneity among a panel of ten YM155-resistant sublines of the neuroblastoma cell line UKF-NB-3. This heterogeneity is of conceptual importance, because it shows that even a defined cancer cell population of limited intrinsic heterogeneity can develop various resistance mechanisms and phenotypes in response to treatment. From a clinical perspective, this means that the close monitoring of cancer cell evolution in response to therapy will have to become an essential part of the design of individualised therapies. Notably, such insights can only be gained from preclinical model systems such as drug-adapted cancer cell lines, which enable the repeated adaptation of a given cancer cell population to the same treatment, but not from clinical material as every patient can only be treated once.

Our findings also demonstrate that biomarkers can indicate resistance formation, even when they do not enable the prediction of drug sensitivity in therapy-naïve cancer cells. Hence, the use of biomarkers differs between the intrinsic and the acquired resistance setting, and pre-clinical models of acquired drug resistance are needed for the identification of such biomarkers that herald resistance development.

## Methods

### Cells

The MYCN-amplified neuroblastoma cell line UKF-NB-3 was established from a bone marrow metastasis of a stage IV neuroblastoma patient [36]. Ten YM155-resistant UKF-NB-3 sublines were derived from the resistant cancer cell line (RCCL) collection (https://research.kent.ac.uk/industrial-biotechnology-centre/the-resistant-cancer-cell-line-rccl-collection/). They were established by adaptation of UKF-NB-3 cells to growth in the presence of YM155 20nM by previously described methods [37] and designated as UKF-NB-3^r^YM155^20nM^I to UKF-NB-3^r^YM155^20nM^X. All cells were propagated in IMDM supplemented with 10 % FBS, 100 IU/ml penicillin and 100 µg/ml streptomycin at 37°C. Cells were routinely tested for mycoplasma contamination and authenticated by short tandem repeat profiling.

To determine doubling times, 2×10^4^ cells per well were plated into 6-well plates, incubated at 37°C and 5% CO_2_, and counted after 1,2,3,5 and 7 days using a Neubauer chamber. Doubling times were then calculated using http://www.doubling-time.com/compute.php.

### Viability assay

Cell viability was tested by the 3-(4,5-dimethylthiazol-2-yl)-2,5-diphenyltetrazolium bromide (MTT) dye reduction assay after 120 h incubation modified as described previously [37].

### *TP53* next generation sequencing

*TP53* next generation sequencing was performed as previously described [15]. All coding exonic and flanking intronic regions of the human TP53 gene were amplified from genomic DNA with Platinum™ Taq DNA polymerase (Life Technologies) by multiplex PCR using two primer pools with 12 non-overlapping primer pairs each, yielding approximately 180 bp amplicons. Each sample was tagged with a unique 8-nucleotide barcode combination using twelve differently barcoded forward and eight differently barcoded reverse primer pools. Barcoded PCR products from up to 96 samples were pooled, purified and an indexed sequencing library was prepared using the NEBNext® ChIP-Seq Library Prep Master Mix Set for Illumina in combination with NEBNext® Multiplex Oligos for Illumina (New England Biolabs). The quality of sequencing libraries was verified on a Bioanalyzer DNA High Sensitivity chip (Agilent) and quantified by digital PCR. 2 × 250 bp paired-end sequencing was carried out on an Illumina MiSeq (Illumina) according to the manufacturer’s recommendations at a mean coverage of 300x.

Read pairs were demultiplexed according to the forward and reverse primers and subsequently aligned using the Burrows-Wheeler Aligner against the Homo sapiens Ensembl reference (rev. 79). Overlapping mate pairs where combined and trimmed to the amplified region. Coverage for each amplicon was calculated via SAMtools (v1.1) [38]. To identify putative mutations, variant calling was performed using SAMtools in combination with VarScan2 (v2.3.9) [39]. Initially, SAMtools was used to create pileups with a base quality filter of 15. Duplicates, orphan reads, unmapped and secondary reads were excluded. Subsequently, Varscan2 was applied to screen for SNVs and InDels separately, using a low-stringency setting with minimal variant frequency of 0.1, a minimum coverage of 20 and a minimum of 10 supporting reads per variant to account for cellular and clonal heterogeneity. Minimum average quality was set to 20 and a strand filter was applied to minimize miscalls due to poor sequencing quality or amplification bias. The resulting list of putative variants was compared against the IARC TP53 (R17) database to check for known p53 cancer mutations.

### Western blot

Cells were lysed using Triton-X-100 sample buffer, and proteins were separated by SDS-PAGE. Detection occurred by using specific antibodies against β-actin (Biovision through BioCat GmbH, Heidelberg Germany), SLC35F2 (Santa Cruz Biotechnology, Dallas, TX, USA), GAPDH, ABCB1 (Cell Signaling via New England Biolabs, Frankfurt, Germany), p53 (Enzo Life Sciences, Lörrach, Germany), and survivin (R&D Systems, Minneapolis, MN, USA). Protein bands were visualized by laser-induced fluorescence using infrared scanner for protein quantification (Odyssey, Li-Cor Biosciences, Lincoln, NE, USA) and Image Studio Ver. 5.2 software (Li-Cor Biosciences) for densitometric analyses.

### RNA interference experiments

Transient depletion of BIRC5/ survivin was achieved using synthetic siRNA oligonucleotides (ON-TARGETplus SMARTpool) from Dharmacon (Lafayette, CO; USA). Non-targeting siRNA (ON-TARGETplus SMARTpool) was used as negative control. Cells were transfected by electroporation using the NEON Transfection System (Invitrogen, Darmstadt; Germany) according to the manufacturer protocol. Cells were grown to 60-80 % confluence, trypsinised, and 1.2 × 10^6^ cells were re-suspended in 200 µl resuspension buffer R including 2.5 µM siRNA. The electroporation was performed using two 20 ms pulses of 1400 V. Subsequently, the cells were transferred into cell culture plates or flasks, containing pre-warmed cell culture medium.

### Irradiation procedure

10^4^ cells per well were irradiated at room temperature in 96 well cell culture plates (Greiner, Bio-ONE GmbH, Frickenhausen, Germany) with single doses of X-rays ranging from 1 to 5 Gy using a linear accelerator (SL 75/5, Elekta, Crawley, UK) with 6 MeV photons/100 cm focus–surface distance with a dose rate of 4.0 Gy/min. Sham-irradiated cultures were kept at room temperature in the X-ray control room while the other samples were irradiated.

### Analysis of data derived from large pharmacogenomic studies

All data (including drug response area under curve (AUC) data for YM-155-treated cancer cell lines, basal gene-expression for ABCB1, BIRC5 (the gene that encodes survivin), and SLC35F2, and genomic alterations of p53) in this study were obtained from two online resources: Version 2 of the Cancer Therapeutics Response Portal (CTRP v2) data [28,40] were obtained from the Cancer Target Discovery and Development (CTD^2^) data portal (ocg.cancer.gov/programs/ctd2/data-portal). The Genomics of Drug Sensitivity in Cancer (GDSC) data was obtained from www.cancerrxgene.org [41,42].

The CTRP contains ABCB1, BIRC5, and SLC35F2 expression data for 823 cell lines and YM-155 AUC data for 715 cell lines. For 703 cell lines (including 12 neuroblastoma cell lines), gene expression data and YM155 AUC values were available. Whole exome sequencing (WES) data was available for 546 of the cell lines for which YM-155 sensitivity data was also available (including 11 neuroblastoma cell lines).

The GDSC contains ABCB1, BIRC5, and SLC35F2 expression data for 1019 cell lines and YM155 AUC data for 945 cell lines. Expression data and WES data were available for all 945 cell lines with YM-155 sensitivity data (including 30 neuroblastoma cell lines).

Data processing was performed using Perl version 5.26.0, and R statistical packages version 3.3.2. Cell lines were determined to display either high or low expression for each gene using the median gene expression as a threshold (i.e. low expression <= median expression, high expression > median expression). Box plots indicating YM-155 sensitivity in cell lines that display low or high expression of a certain gene or wild-type or mutant *TP53* were produced using the ggplot2 package [43] in R.

Statistical tests were carried out in R and included Wilcoxon rank-sum test [44] and Pearson’s correlation [45]. Correction for multiple comparisons was performed using the Benjamini-Hochberg procedure [46].

### Statistics

Results are expressed as mean ± S.D. of at least three experiments. Comparisons between two groups were performed using Student’s t-test. Three or more groups were compared by ANOVA followed by the Student-Newman-Keuls test. P values lower than 0.05 were considered to be significant.

## References

1. Nakahara T, Kita A, Yamanaka K, Mori M, Amino N, Takeuchi M, Tominaga F, Hatakeyama S, Kinoyama I, Matsuhisa A, Kudoh M, Sasamata M. YM155, a novel small-molecule survivin suppressant, induces regression of established human hormone-refractory prostate tumor xenografts. Cancer Res. 2007;67:8014–21.

2. Tang H, Shao H, Yu C, Hou J. Mcl-1 downregulation by YM155 contributes to its synergistic anti-tumor activities with ABT-263. Biochem Pharmacol. 2011;82:1066–72.

3. Glaros TG, Stockwin LH, Mullendore ME, Smith B, Morrison BL, Newton DL. The “survivin suppressants” NSC 80467 and YM155 induce a DNA damage response. Cancer Chemother Pharmacol. 2012;70:207–12.

4. Na YS, Yang SJ, Kim SM, Jung KA, Moon JH, Shin JS, Yoon DH, Hong YS, Ryu MH, Lee JL, Lee JS, Kim TW. YM155 induces EGFR suppression in pancreatic cancer cells. PLoS One. 2012;7:e38625.

5. Rauch A, Hennig D, Schäfer C, Wirth M, Marx C, Heinzel T, Schneider G, Krämer OH. Survivin and YM155: how faithful is the liaison? Biochim Biophys Acta. 2014; 1845: 202–20.

6. Cheng SM, Chang YC, Liu CY, Lee JY, Chan HH, Kuo CW, Lin KY, Tsai SL, Chen SH, Li CF, Leung E, Kanwar JR, Huang CC, Chang JY, Cheung CH. YM155 down-regulates survivin and XIAP, modulates autophagy and induces autophagy-dependent DNA damage in breast cancer cells. Br J Pharmacol. 2015;172:214–34.

7. de Necochea-Campion R, Diaz Osterman CJ, Hsu HW, Fan J, Mirshahidi S, Wall NR, Chen CS. AML sensitivity to YM155 is modulated through AKT and Mcl-1. Cancer Lett. 2015; 366: 44–51.

8. Jung SA, Park YM, Hong SW, Moon JH, Shin JS, Lee HR, Ha SH, Lee DH, Kim JH, Kim SM, Kim JE, Kim KP, Hong YS, Choi EK, Lee JS, Jin DH, Kim T. Cellular inhibitor of apoptosis protein 1 (cIAP1) stability contributes to YM155 resistance in human gastric cancer cells. J Biol Chem. 2015;290:9974–85.

9. Pennati M, Sbarra S, De Cesare M, Lopergolo A, Locatelli SL, Campi E, Daidone MG, Carlo-Stella C, Gianni AM, Zaffaroni N. YM155 sensitizes triple-negative breast cancer to membrane-bound TRAIL through p38 MAPK- and CHOP-mediated DR5 upregulation. Int J Cancer. 2015;136:299–309.

10. Zhao X, Puszyk WM, Lu Z, Ostrov DA, George TJ, Robertson KD, Liu C. Small molecule inhibitor YM155-mediated activation of death receptor 5 is crucial for chemotherapy-induced apoptosis in pancreatic carcinoma. Mol Cancer Ther. 2015;14:80–9.

11. Ho SH, Ali A, Chin TM, Go ML. Dioxonaphthoimidazoliums AB1 and YM155 disrupt phosphorylation of p50 in the NF-κB pathway. Oncotarget. 2016;7:11625–36.

12. Kojima Y, Hayakawa F, Morishita T, Sugimoto K, Minamikawa Y, Iwase M, Yamamoto H, Hirano D, Imoto N, Shimada K, Okada S, Kiyoi H. YM155 induces apoptosis through proteasome-dependent degradation of MCL-1 in primary effusion lymphoma. Pharmacol Res. 2017;120:242–51.

13. Lamers F, Schild L, Koster J, Versteeg R, Caron HN, Molenaar JJ. Targeted BIRC5 silencing using YM155 causes cell death in neuroblastoma cells with low ABCB1 expression. Eur J Cancer. 2012;48:763–71.

14. Liang H, Zhang L, Xu R, Ju XL. Silencing of survivin using YM155 induces apoptosis and chemosensitization in neuroblastomas cells. Eur Rev Med Pharmacol Sci. 2013;17:2909–15.

15. Voges Y, Michaelis M, Rothweiler F, Schaller T, Schneider C, Politt K, Mernberger M, Nist A, Stiewe T, Wass MN, Rödel F, Cinatl J Jr. Effects of YM155 on survivin levels and viability in neuroblastoma cells with acquired drug resistance. Cell Death Dis. 2016;7:e2410.

16. Radic-Sarikas B, Halasz M, Huber KVM, Winter GE, Tsafou KP, Papamarkou T, Brunak S, Kolch W, Superti-Furga G. Lapatinib potentiates cytotoxicity of YM155 in neuroblastoma via inhibition of the ABCB1 efflux transporter. Sci Rep. 2017;7:3091.

17. Pinto NR, Applebaum MA, Volchenboum SL, Matthay KK, London WB, Ambros PF, Nakagawara A, Berthold F, Schleiermacher G, Park JR, Valteau-Couanet D, Pearson AD, Cohn SL. Advances in Risk Classification and Treatment Strategies for Neuroblastoma. J Clin Oncol. 2015;33:3008–17.

18. Bagatell R, Cohn SL. Genetic discoveries and treatment advances in neuroblastoma. Curr Opin Pediatr. 2016;28:19–25.

19. Speleman F, Park JR, Henderson TO. Neuroblastoma: A Tough Nut to Crack. Am Soc Clin Oncol Educ Book. 2016;35:e548–57.

20. PDQ Pediatric Treatment Editorial Board. Neuroblastoma Treatment (PDQ®): Health Professional Version. PDQ Cancer Information Summaries [Internet]. Bethesda (MD): National Cancer Institute (US); 2002-. 2017 Apr 14.

21. Moreno L, Caron H, Geoerger B, Eggert A, Schleiermacher G, Brock P, Valteau-Couanet D, Chesler L, Schulte JH, De Preter K, Molenaar J, Schramm A, Eilers M, Van Maerken T, Johnsen JI, Garrett M, George SL, Tweddle DA, Kogner P, Berthold F, Koster J, Barone G, Tucker ER, Marshall L, Herold R, Sterba J, Norga K, Vassal G, Pearson AD. Accelerating Drug Development for Neuroblastoma - New Drug Development Strategy: An Innovative Therapies for Children with Cancer, European Network for Cancer Research in Children and Adolescents and International Society of Paediatric Oncology Europe Neuroblastoma Project. Expert Opin Drug Discov. 2017;12:801–11.

22. Holohan C, Van Schaeybroeck S, Longley DB, Johnston PG. Cancer drug resistance: an evolving paradigm. Nat Rev Cancer. 2013;13:714–26.

23. Perakis S, Speicher MR. Emerging concepts in liquid biopsies. BMC Med. 2017;15:75.

24. Miklos W, Pelivan K, Kowol CR, Pirker C, Dornetshuber-Fleiss R, Spitzwieser M, Englinger B, van Schoonhoven S, Cichna-Markl M, Koellensperger G, Keppler BK, Berger W, Heffeter P. Triapine-mediated ABCB1 induction via PKC induces widespread therapy unresponsiveness but is not underlying acquired triapine resistance. Cancer Lett. 2015;361:112–20.

25. Hata AN, Niederst MJ, Archibald HL, Gomez-Caraballo M, Siddiqui FM, Mulvey HE, Maruvka YE, Ji F, Bhang HE, Krishnamurthy Radhakrishna V, Siravegna G, Hu H, Raoof S, Lockerman E, Kalsy A, Lee D, Keating CL, Ruddy DA, Damon LJ, Crystal AS, Costa C, Piotrowska Z, Bardelli A, Iafrate AJ, Sadreyev RI, Stegmeier F, Getz G, Sequist LV, Faber AC, Engelman JA. Tumor cells can follow distinct evolutionary paths to become resistant to epidermal growth factor receptor inhibition. Nat Med. 2016;22:262–9.

26. Carter L, Rothwell DG, Mesquita B, Smowton C, Leong HS, Fernandez-Gutierrez F, Li Y, Burt DJ, Antonello J, Morrow CJ, Hodgkinson CL, Morris K, Priest L, Carter M, Miller C, Hughes A, Blackhall F, Dive C, Brady G. Molecular analysis of circulating tumor cells identifies distinct copy-number profiles in patients with chemosensitive and chemorefractory small-cell lung cancer. Nat Med. 2017;23:114–9.

27. Lipinski KA, Barber LJ, Davies MN, Ashenden M, Sottoriva A, Gerlinger M. Cancer Evolution and the Limits of Predictability in Precision Cancer Medicine. Trends Cancer. 2016;2:49–63.

28. Basu A, Bodycombe NE, Cheah JH, Price EV, Liu K, Schaefer GI, Ebright RY, Stewart ML, Ito D, Wang S, Bracha AL, Liefeld T, Wawer M, Gilbert JC, Wilson AJ, Stransky N, Kryukov GV, Dancik V, Barretina J, Garraway LA, Hon CS, Munoz B, Bittker JA, Stockwell BR, Khabele D, Stern AM, Clemons PA, Shamji AF, Schreiber SL. An interactive resource to identify cancer genetic and lineage dependencies targeted by small molecules. Cell. 2013;154:1151–61.

29. Garnett MJ, Edelman EJ, Heidorn SJ, Greenman CD, Dastur A, Lau KW, Greninger P, Thompson IR, Luo X, Soares J, Liu Q, Iorio F, Surdez D, Chen L, Milano RJ, Bignell GR, Tam AT, Davies H, Stevenson JA, Barthorpe S, Lutz SR, Kogera F, Lawrence K, McLaren-Douglas A, Mitropoulos X, Mironenko T, Thi H, Richardson L, Zhou W, Jewitt F, Zhang T, O’Brien P, Boisvert JL, Price S, Hur W, Yang W, Deng X, Butler A, Choi HG, Chang JW, Baselga J, Stamenkovic I, Engelman JA, Sharma SV, Delattre O, Saez-Rodriguez J, Gray NS, Settleman J, Futreal PA, Haber DA, Stratton MR, Ramaswamy S, McDermott U, Benes CH. Systematic identification of genomic markers of drug sensitivity in cancer cells. Nature. 2012;483:570–5.

30. Nakahara T, Kita A, Yamanaka K, Mori M, Amino N, Takeuchi M, Tominaga F, Kinoyama I, Matsuhisa A, Kudou M, Sasamata M. Broad spectrum and potent antitumor activities of YM155, a novel small-molecule survivin suppressant, in a wide variety of human cancer cell lines and xenograft models. Cancer Sci. 2011;102:614–21.

31. Ghadimi MP, Young ED, Belousov R, Zhang Y, Lopez G, Lusby K, Kivlin C, Demicco EG, Creighton CJ, Lazar AJ, Pollock RE, Lev D. Survivin is a viable target for the treatment of malignant peripheral nerve sheath tumors. Clin Cancer Res. 2012;18:2545–57.

32. Xia H, Chen J, Shi M, Deivasigamani A, Ooi LL, Hui KM. The over-expression of survivin enhances the chemotherapeutic efficacy of YM155 in human hepatocellular carcinoma. Oncotarget. 2015;6:5990–6000.

33. Sim MY, Huynh H, Go ML, Yuen JSP. Action of YM155 on clear cell renal cell carcinoma does not depend on survivin expression levels. PLoS One. 2017;12:e0178168.

34. Winter GE, Radic B, Mayor-Ruiz C, Blomen VA, Trefzer C, Kandasamy RK, Huber KVM, Gridling M, Chen D, Klampfl T, Kralovics R, Kubicek S, Fernandez-Capetillo O, Brummelkamp TR, Superti-Furga G. The solute carrier SLC35F2 enables YM155-mediated DNA damage toxicity. Nat Chem Biol. 2014;10:768–73.

35. Chang BH, Johnson K, LaTocha D, Rowley JS, Bryant J, Burke R, Smith RL, Loriaux M, Müschen M, Mullighan C, Druker BJ, Tyner JW. YM155 potently kills acute lymphoblastic leukemia cells through activation of the DNA damage pathway. J Hematol Oncol. 2015;8:39.

36. Kotchetkov R, Driever PH, Cinatl J, Michaelis M, Karaskova J, Blaheta R, Squire JA, Von Deimling A, Moog J, Cinatl J Jr. Increased malignant behavior in neuroblastoma cells with acquired multi-drug resistance does not depend on P-gp expression. Int J Oncol. 2005;27:1029–37.

37. Michaelis M, Rothweiler F, Barth S, Cinatl J, van Rikxoort M, Löschmann N, Voges Y, Breitling R, von Deimling A, Rödel F, Weber K, Fehse B, Mack E, Stiewe T, Doerr HW, Speidel D, Cinatl J Jr. Adaptation of cancer cells from different entities to the MDM2 inhibitor nutlin-3 results in the emergence of p53-mutated multi-drug-resistant cancer cells. Cell Death Dis. 2011;2:e243.

38. Li H, Handsaker B, Wysoker A, Fennell T, Ruan J, Homer N, Marth G, Abecasis G, Durbin R; 1000 Genome Project Data Processing Subgroup. The Sequence Alignment/Map format and SAMtools. Bioinformatics. 2009;25:2078–9.

39. Koboldt DC, Zhang Q, Larson DE, Shen D, McLellan MD, Lin L, Miller CA, Mardis ER, Ding L, Wilson RK. VarScan 2: somatic mutation and copy number alteration discovery in cancer by exome sequencing. Genome Res. 2012;22:568–76.

40. Rees MG, Seashore-Ludlow B, Cheah JH, Adams DJ, Price EV, Gill S, Javaid S, Coletti ME, Jones VL, Bodycombe NE, Soule CK, Alexander B, Li A, Montgomery P, Kotz JD, Hon CS, Munoz B, Liefeld T, Dancík V, Haber DA, Clish CB, Bittker JA, Palmer M, Wagner BK, Clemons PA, Shamji AF, Schreiber SL. Correlating chemical sensitivity and basal gene expression reveals mechanism of action. Nat Chem Biol. 2016;12:109–16.

41. Yang W, Soares J, Greninger P, Edelman EJ, Lightfoot H, Forbes S, Bindal N, Beare D, Smith JA, Thompson IR, Ramaswamy S, Futreal PA, Haber DA, Stratton MR, Benes C, McDermott U, Garnett MJ. Genomics of Drug Sensitivity in Cancer (GDSC): a resource for therapeutic biomarker discovery in cancer cells. Nucleic Acids Res. 2013;41:D955–61.

42. Iorio F, Knijnenburg TA, Vis DJ, Bignell GR, Menden MP, Schubert M, Aben N, Gonçalves E, Barthorpe S, Lightfoot H, Cokelaer T, Greninger P, van Dyk E, Chang H, de Silva H, Heyn H, Deng X, Egan RK, Liu Q, Mironenko T, Mitropoulos X, Richardson L, Wang J, Zhang T, Moran S, Sayols S, Soleimani M, Tamborero D, Lopez-Bigas N, Ross-Macdonald P, Esteller M, Gray NS, Haber DA, Stratton MR, Benes CH, Wessels LF, Saez-Rodriguez J, McDermott U, Garnett MJ. A Landscape of Pharmacogenomic Interactions in Cancer. Cell. 2016;166:740–54.

43. Wickham, H. ggplot2: Elegant Graphics for Data Analysis. Berlin, Germany: Springer, 2009.

44. Mann HB, Whitney DR. On a Test of Whether one of Two Random Variables is Stochastically Larger than the Other. Ann Math Stat. 1947;18:50–60.

45. Pearson K. Notes on regression and inheritance in the case of two parents. Proc R Soc Lond. 1895;58:240–2.

46. Benjamin Y, Hochberg Y. “Controlling the false discovery rate: a practical and powerful approach to multiple testing”. J R Stat Soc B. 1995;57:289–300.

