## Supplementary material for "Drug-adapted cancer cell lines reveal drug-induced heterogeneity and enable the identification of biomarker candidates for the acquired resistance setting": Suppl Files

**Table S1**. YM155 concentrations (nM) that reduce the viability of UKF-NB-3 or YM155-adapted UKF-NB-3 sub-lines by 50% ($IC_{50}$) or 90% ($IC_{90}$) as indicated by MTT assay after 120h of incubation and doubling times of the cells.

| | $IC_{50}$ | $IC_{90}$ | Doubling times (h) |
|---|---|---|---|
| UKF-NB-3 | 0.55 ± 0.06 | 1.01 ± 0.24 | 30.8 ± 0.8 |
| UKF-NB-3rYM155$^{20nM}$I | 36.2 ± 2.0 (66)[1] | 94.8 ± 0.7 (94)[2] | 41.5 ± 5.2 |
| UKF-NB-3rYM155$^{20nM}$II | 23.6 ± 2.2 (43) | 39.9 ± 0.9 (40) | 39.2 ± 4.8 |
| UKF-NB-3rYM155$^{20nM}$III | 31.8 ± 2.3 (58) | 49.1 ± 0.2 (49) | 32.3 ± 1.1 |
| UKF-NB-3rYM155$^{20nM}$IV | 21.0 ± 0.6 (38) | 29.8 ± 4.8 (30) | 32.2 ± 2.0 |
| UKF-NB-3rYM155$^{20nM}$V | 25.1 ± 0.1 (45) | 39.5 ± 0.7 (39) | 35.3 ± 2.8 |
| UKF-NB-3rYM155$^{20nM}$VI | 41.9 ± 5.3 (76) | 136 ± 7 (135) | 36.3 ± 2.2 |
| UKF-NB-3rYM155$^{20nM}$VII | 36.5 ± 5.5 (66) | 96.2 ± 23.9 (95) | 46.1 ± 2.1 |
| UKF-NB-3rYM155$^{20nM}$VIII | 34.5 ± 0.6 (63) | 84.7 ± 27.3 (84) | 47.0 ± 2.5 |
| UKF-NB-3rYM155$^{20nM}$IX | 27.2 ± 0.5 (49) | 44.8 ± 2.9 (44) | 33.3 ± 0.2 |
| UKF-NB-3rYM155$^{20nM}$X | 26.9 ± 0.5 (49) | 45.2 ± 4.7 (45) | 41.9 ± 2.2 |

[1] fold resistance ($IC_{50}$ YM155-adapted sub-line/ $IC_{50}$ UKF-NB-3)
[2] fold resistance ($IC_{90}$ YM155-adapted sub-line/ $IC_{90}$ UKF-NB-3)

**Table S2**. Drug concentrations that reduce the viability of UKF-NB-3 or YM155-adapted UKF-NB-3 sub-lines by 50% ($IC_{50}$) as indicated by MTT assay after 120h of incubation.

| | Nutlin-3 | Vincristine | Cisplatin | Gemcitabine | Topotecan |
|---|---|---|---|---|---|
| | $IC_{50}$ (µM) | $IC_{50}$ (ng/mL) | $IC_{50}$ (ng/mL) | $IC_{50}$ (ng/mL) | $IC_{50}$ (ng/mL) |
| UKF-NB-3 | 1.05 ± 0.25 | 1.75 ± 0.55 | 169 ± 29 | 0.30 ± 0.03 | 1.29 ± 0.52 |
| UKF-NB-3rYM155$^{20nM}$I | 0.57 ± 0.07 (0.5)[1] | 45.5 ± 11.1 (26) | 157 ± 54 (0.9) | 0.64 ± 0.02 (2.1) | 1.37 ± 0.53 (1.1) |
| UKF-NB-3rYM155$^{20nM}$II | 1.31 ± 0.03 (1.2) | 27.0 ± 12.6 (15) | 183 ± 51 (1.1) | 0.50 ± 0.04 (1.7) | 1.25 ± 0.53 (1.0) |
| UKF-NB-3rYM155$^{20nM}$III | 1.27 ± 0.01 (1.2) | 10.8 ± 6.4 (6.2) | 122 ± 24 (0.7) | 0.62 ± 0.01 (2.1) | 1.06 ± 0.24 (0.8) |
| UKF-NB-3rYM155$^{20nM}$IV | 0.47 ± 0.03 (0.4) | 18.5 ± 8.4 (11) | 159 ± 38 (0.9) | 0.23 ± 0.04 (0.8) | 1.56 ± 0.65 (1.2) |
| UKF-NB-3rYM155$^{20nM}$V | 0.99 ± 0.13 (0.9) | 8.90 ± 7.39 (5.1) | 156 ± 84 (0.9) | 0.12 ± 0.04 (0.4) | 0.91 ± 0.41 (0.7) |
| UKF-NB-3rYM155$^{20nM}$VI | 0.64 ± 0.01 (0.6) | 714 ± 456 (408) | 132 ± 39 (0.8) | 0.64 ± 0.01 (2.1) | 1.55 ± 0.72 (1.2) |
| UKF-NB-3rYM155$^{20nM}$VII | 1.27 ± 0.04 (1.2) | 28.8 ± 10.2 (16) | 134 ± 6 (0.8) | 0.19 ± 0.01 (0.6) | 1.44 ± 0.84 (1.1) |
| UKF-NB-3rYM155$^{20nM}$VIII | 0.70 ± 0.01 (0.7) | 39.5 ± 15.4 (23) | 190 ± 56 (1.1) | 0.65 ± 0.01 (2.2) | 1.26 ± 0.50 (1.0) |
| UKF-NB-3rYM155$^{20nM}$IX | 0.33 ± 0.01 (0.3) | 5.63 ± 1.94 (3.2) | 178 ± 41 (1.1) | 0.18 ± 0.01 (0.6) | 1.59 ± 0.74 (1.2) |
| UKF-NB-3rYM155$^{20nM}$X | 0.64 ± 0.15 (0.6) | 26.0 ± 6.2 (15) | 144 ± 44 (0.9) | 0.54 ± 0.01 (1.8) | 1.21 ± 0.40 (0.9) |

[1] fold resistance ($IC_{50}$ YM155-adapted sub-line/ $IC_{50}$ UKF-NB-3)

**Figure S1**

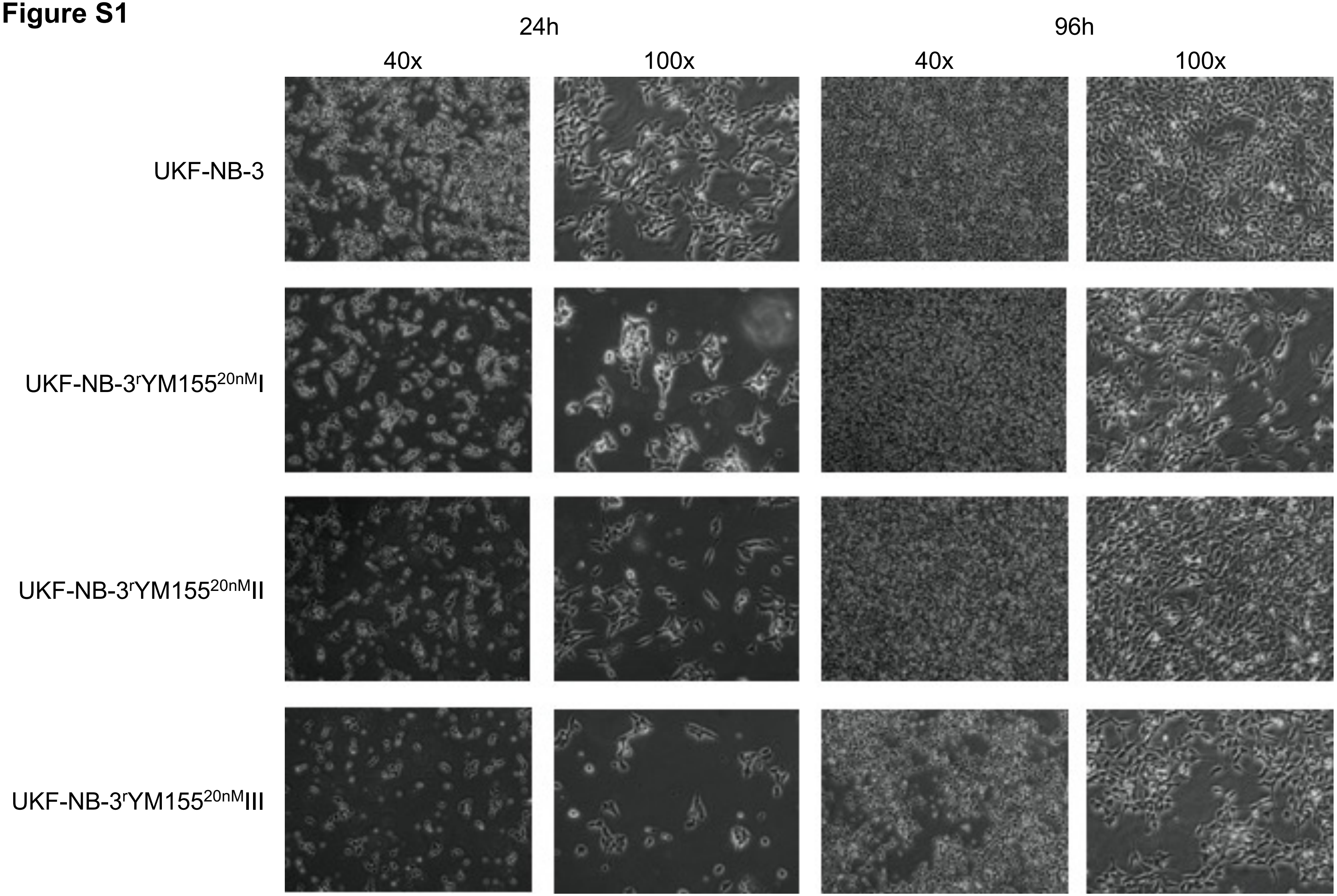
24h
96h
40x
100x
40x
100x
UKF-NB-3
UKF-NB-3$^{r}$YM155$^{20nM}$I
UKF-NB-3$^{r}$YM155$^{20nM}$II
UKF-NB-3$^{r}$YM155$^{20nM}$III

**Figure S1**

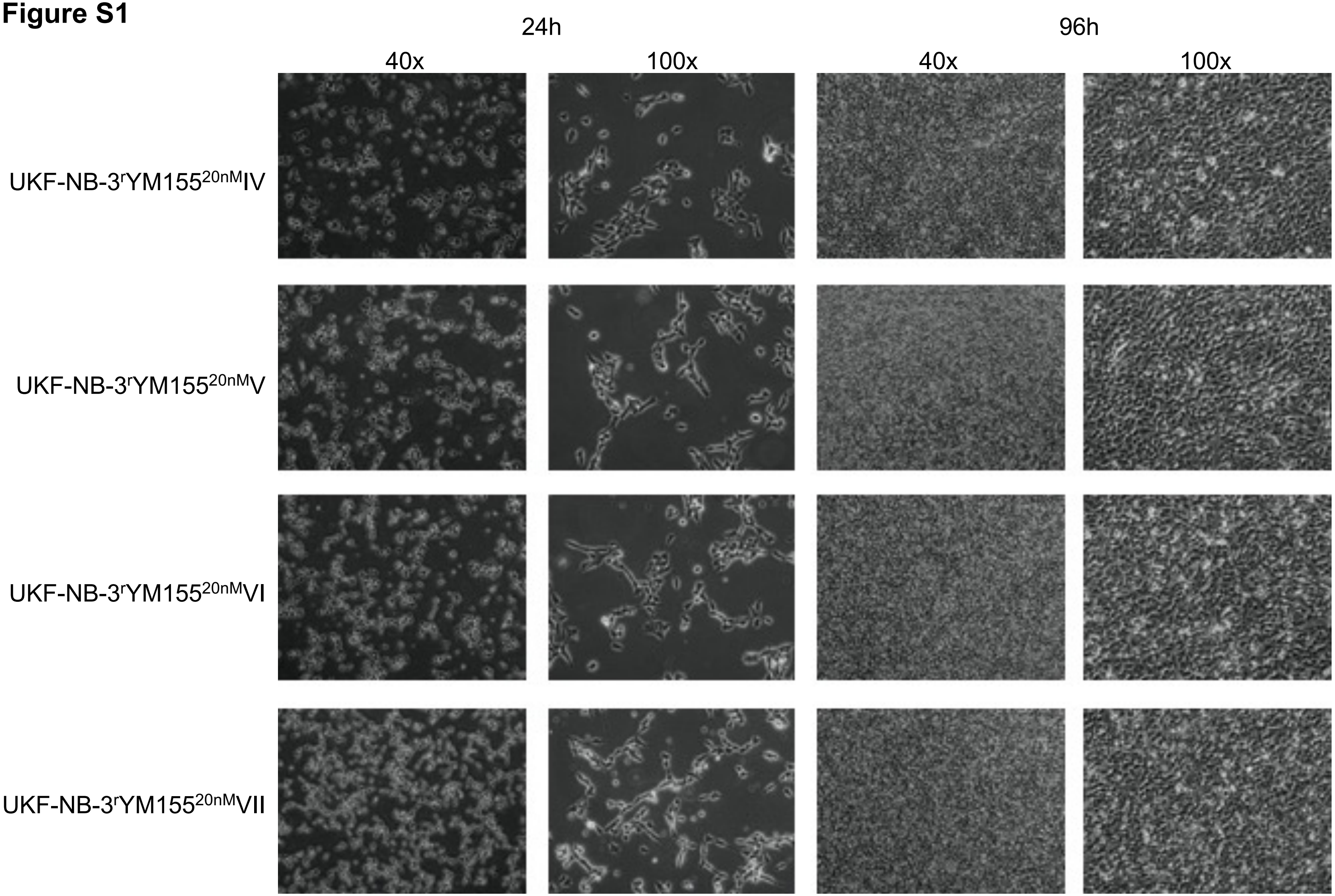
24h
96h
40x
100x
40x
100x
UKF-NB-3rYM155 20nM IV
UKF-NB-3rYM155 20nM V
UKF-NB-3rYM155 20nM VI
UKF-NB-3rYM155 20nM VII

**Figure S1**

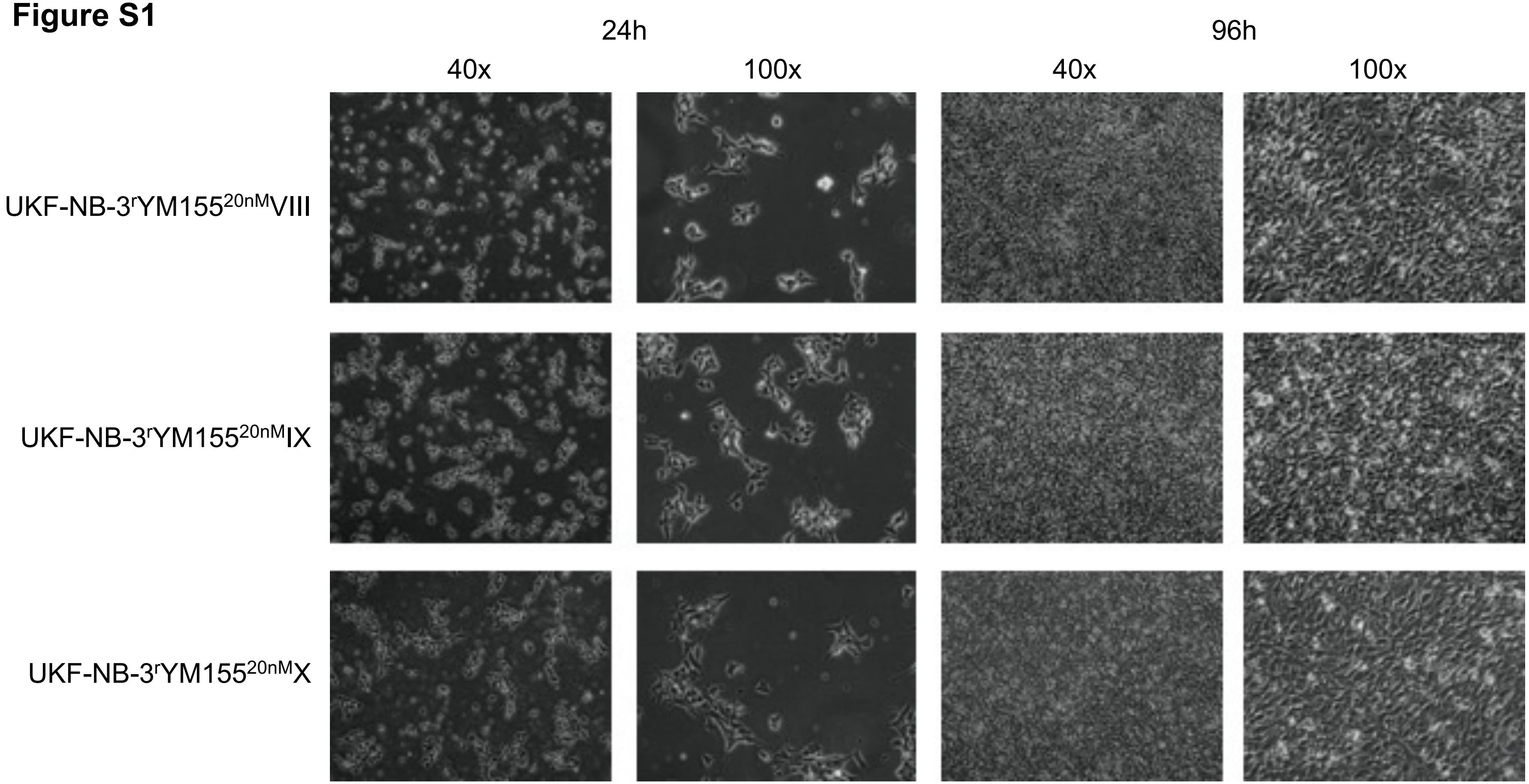


**Figure S1**. Representative photos of the project cell lines indicating cell morphology after different periods of cultivation and at different maginifications.

#### Figure S2

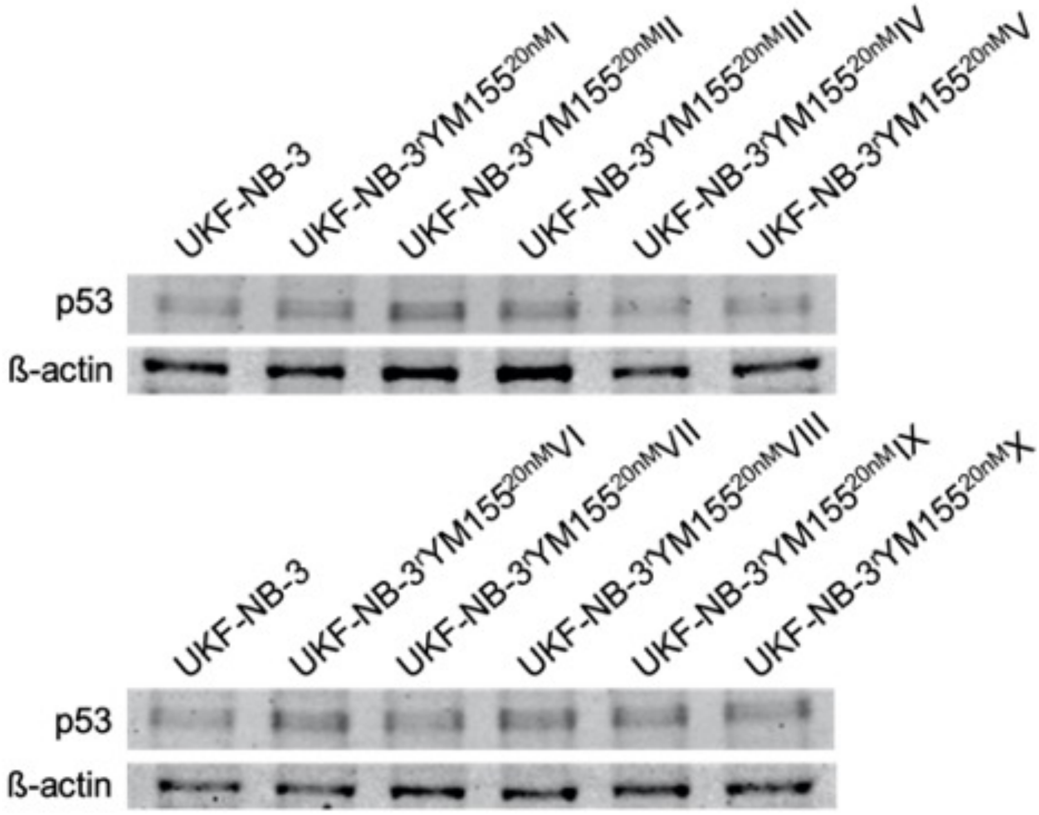


**Figure S2**. p53 levels in UKF-NB-3 and its YM155-adapted sublines.

**Figure S2**. Representative Western blots indicating cellular levels of p53 in UKF-NB-3 and YM155-adapted UKF-NB-3 sub-lines. Densitometric analysis was performed with QuantiOne (BioRad). p53 levels were normalised to β-actin expression and values relative to control cells are displayed.

**Figure S2**

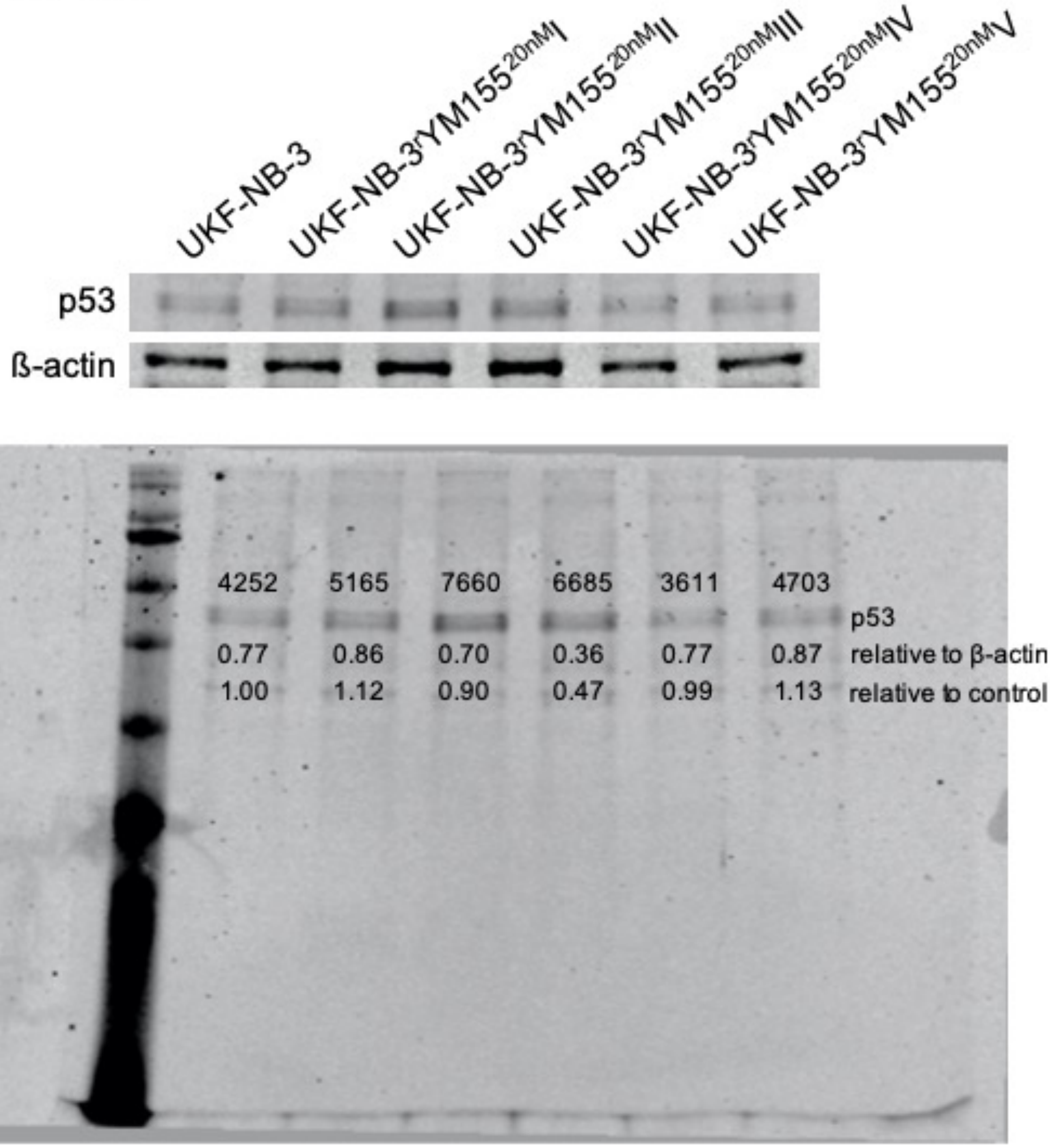

UKF-NB-3
UKF-NB-3rYM155 20nM I
UKF-NB-3rYM155 20nM II
UKF-NB-3rYM155 20nM III
UKF-NB-3rYM155 20nM IV
UKF-NB-3rYM155 20nM V
p53
ß-actin
4252 5165 7660 6685 3611 4703
p53
0.77 0.86 0.70 0.36 0.77 0.87 relative to β-actin
1.00 1.12 0.90 0.47 0.99 1.13 relative to control

**Figure S2**

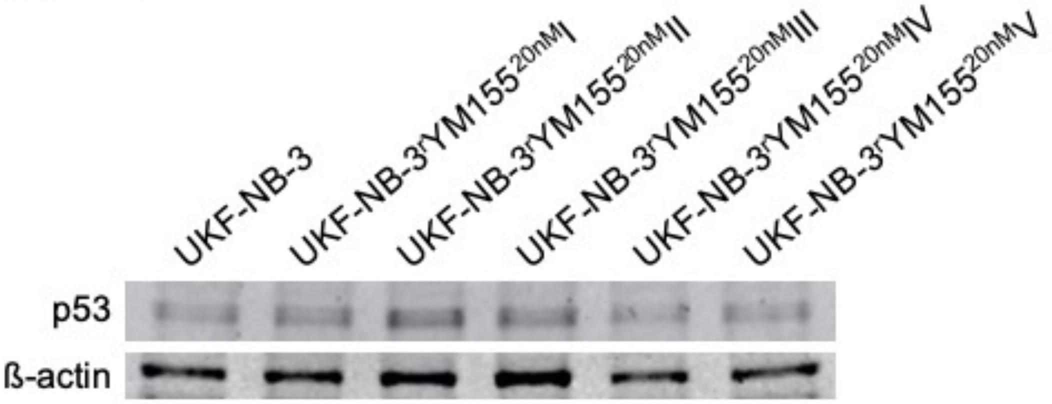
UKF-NB-3
UKF-NB-3rYM155 20nM I
UKF-NB-3rYM155 20nM II
UKF-NB-3rYM155 20nM III
UKF-NB-3rYM155 20nM IV
UKF-NB-3rYM155 20nM V
p53
ß-actin

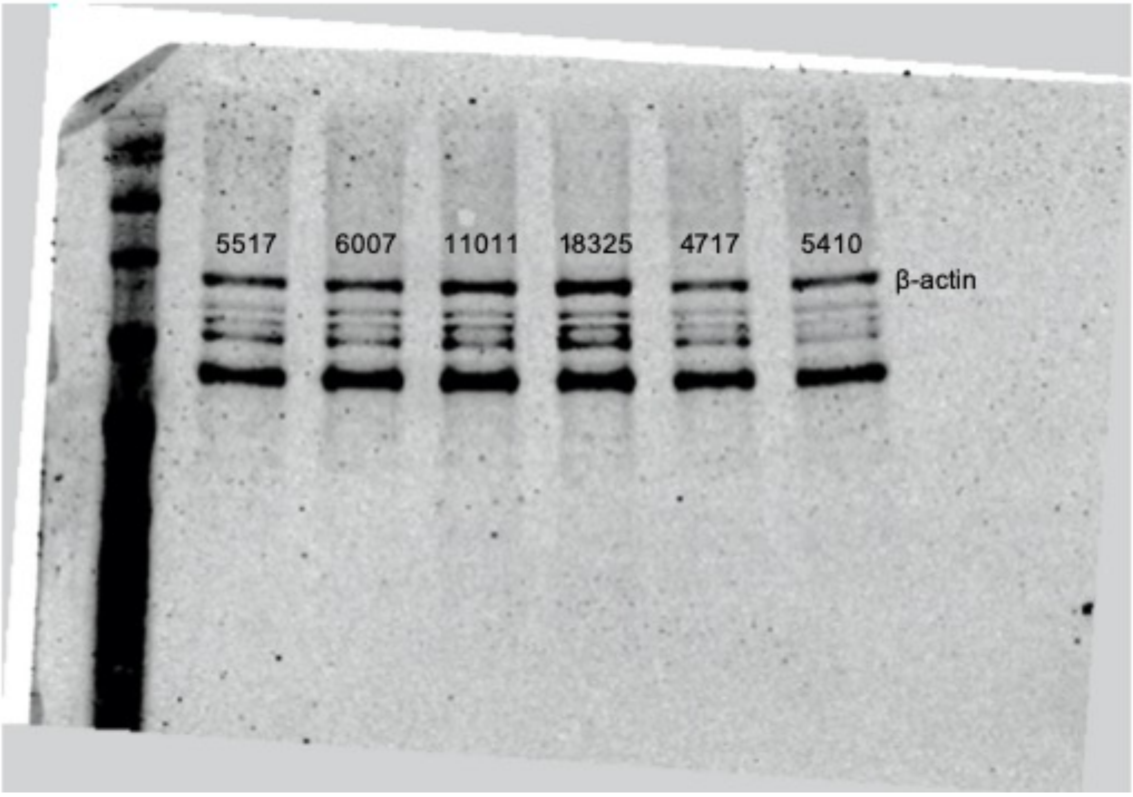
5517
6007
11011
18325
4717
5410
β-actin

**Figure S2**

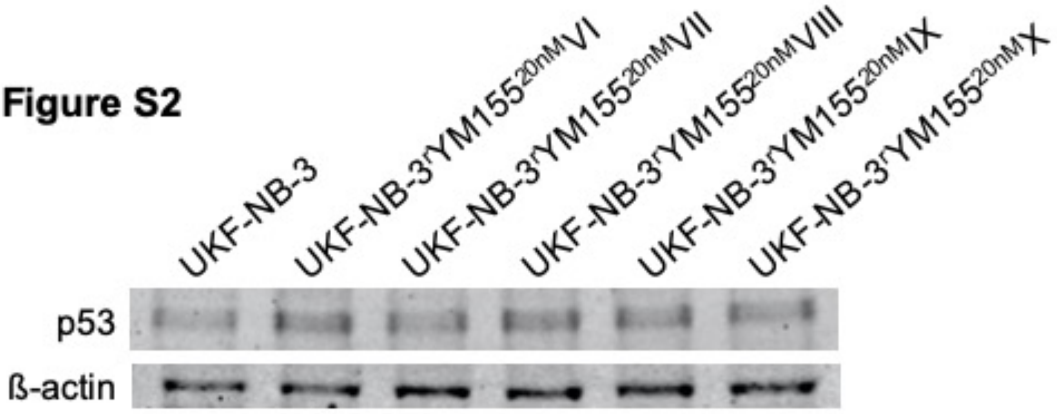
UKF-NB-3
UKF-NB-3$^{r}$YM155$^{20nM}$VI
UKF-NB-3$^{r}$YM155$^{20nM}$VII
UKF-NB-3$^{r}$YM155$^{20nM}$VIII
UKF-NB-3$^{r}$YM155$^{20nM}$IX
UKF-NB-3$^{r}$YM155$^{20nM}$X
p53
ß-actin

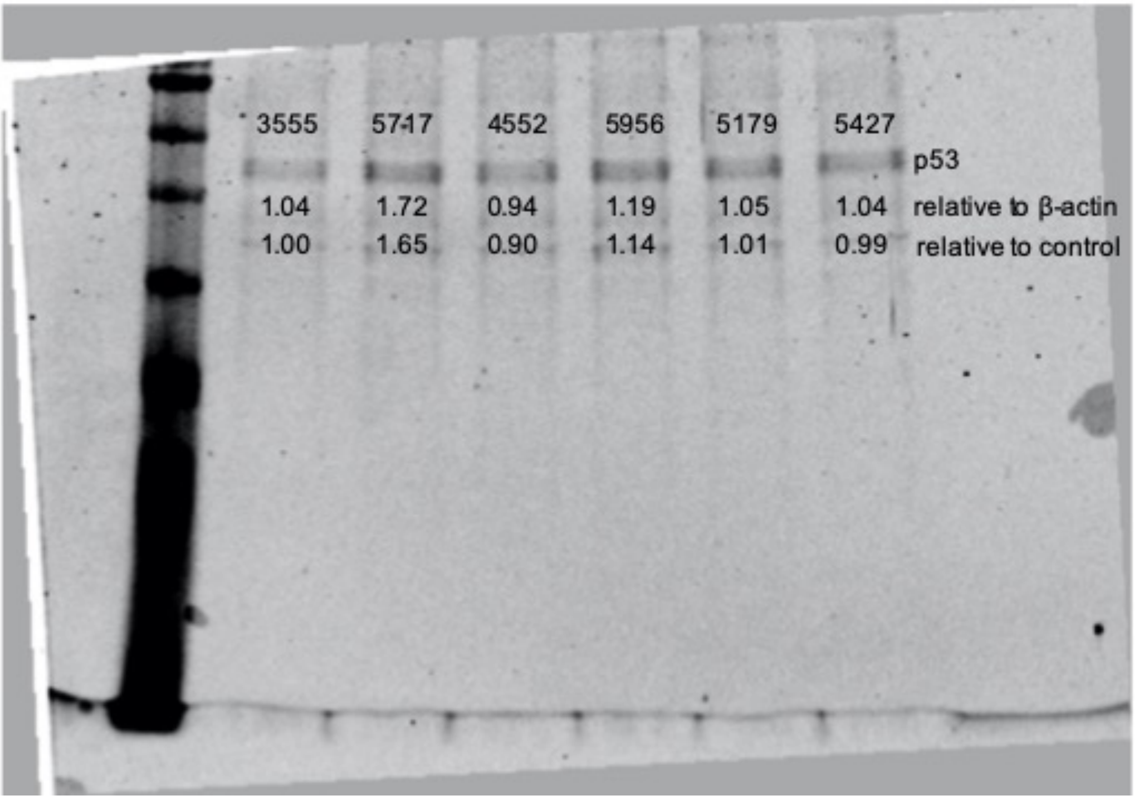
3555 5717 4552 5956 5179 5427
p53
1.04 1.72 0.94 1.19 1.05 1.04 relative to β-actin
1.00 1.65 0.90 1.14 1.01 0.99 relative to control

### Figure S2

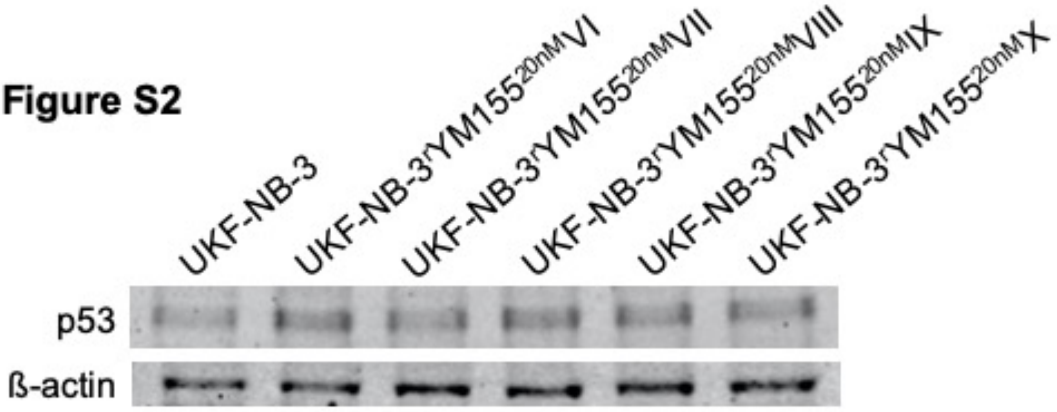

UKF-NB-3
UKF-NB-3$^{r}$YM155$^{20nM}$VI
UKF-NB-3$^{r}$YM155$^{20nM}$VII
UKF-NB-3$^{r}$YM155$^{20nM}$VIII
UKF-NB-3$^{r}$YM155$^{20nM}$IX
UKF-NB-3$^{r}$YM155$^{20nM}$X
p53
ß-actin


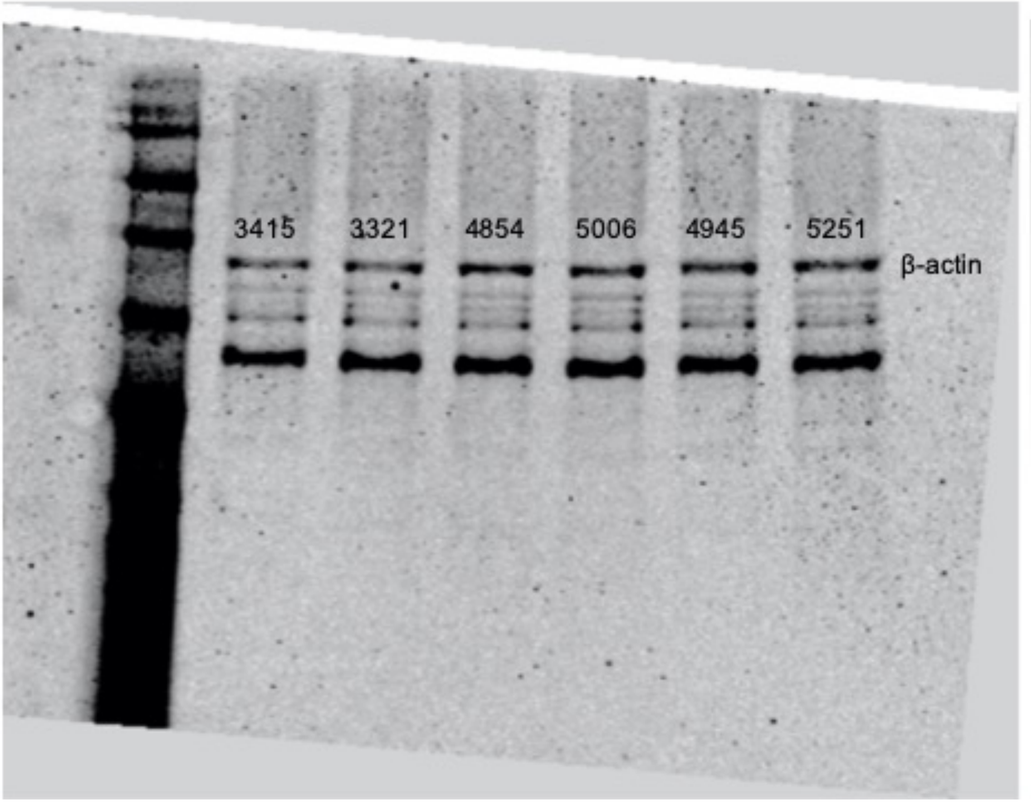

3415
3321
4854
5006
4945
5251
β-actin

#### Figure S3

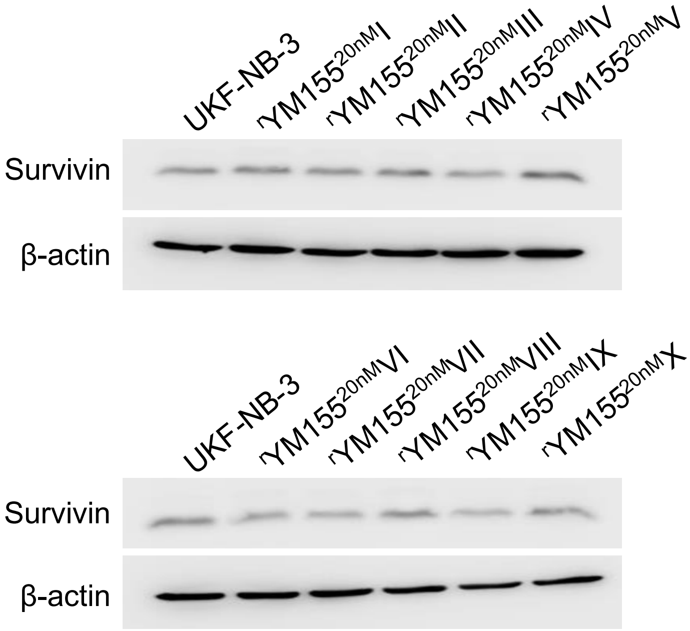


**Figure S3**. Representative Western blots indicating cellular levels of survivin in UKF-NB-3 and YM155-adapted UKF-NB-3 sub-lines. Densitometric analysis was performed with Image Studio Ver. 5.2 software (LICOR). Survivin levels were normalised to β-actin expression and values relative to control cells are displayed.

### Figure S3

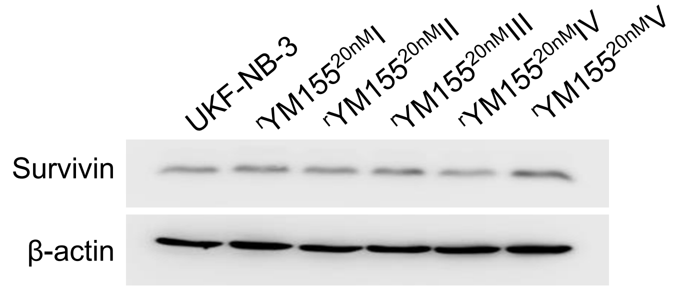
UKF-NB-3
rYM155 20nM I
rYM155 20nM II
rYM155 20nM III
rYM155 20nM IV
rYM155 20nM V
Survivin
β-actin

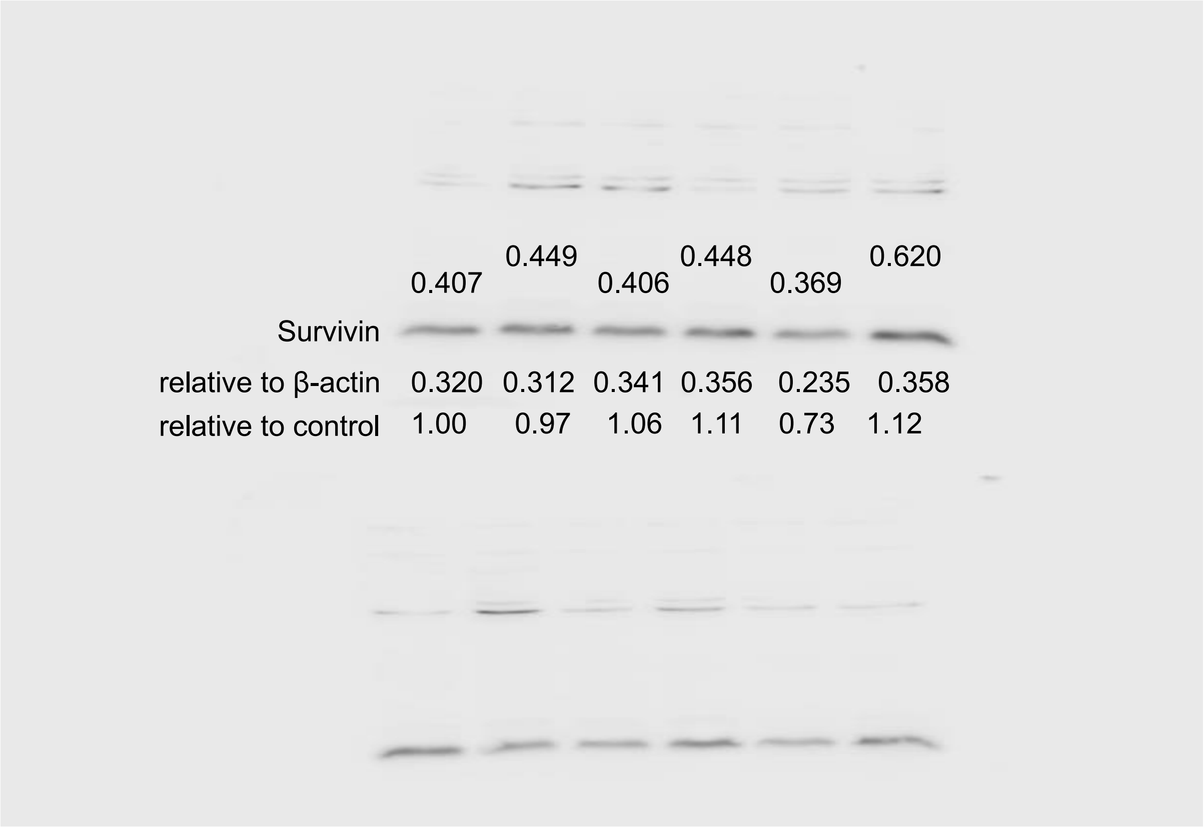
0.407
0.449
0.406
0.448
0.369
0.620
Survivin
relative to β-actin 0.320 0.312 0.341 0.356 0.235 0.358
relative to control 1.00 0.97 1.06 1.11 0.73 1.12

### Figure S3

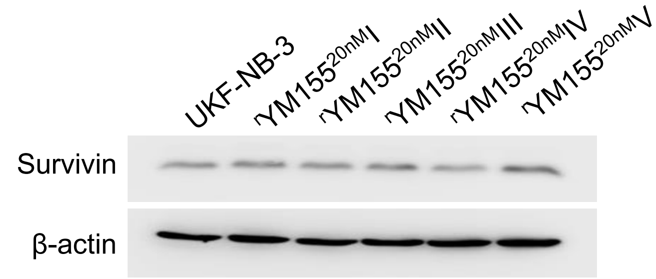
UKF-NB-3
rYM155 20nM I
rYM155 20nM II
rYM155 20nM III
rYM155 20nM IV
rYM155 20nM V
Survivin
β-actin

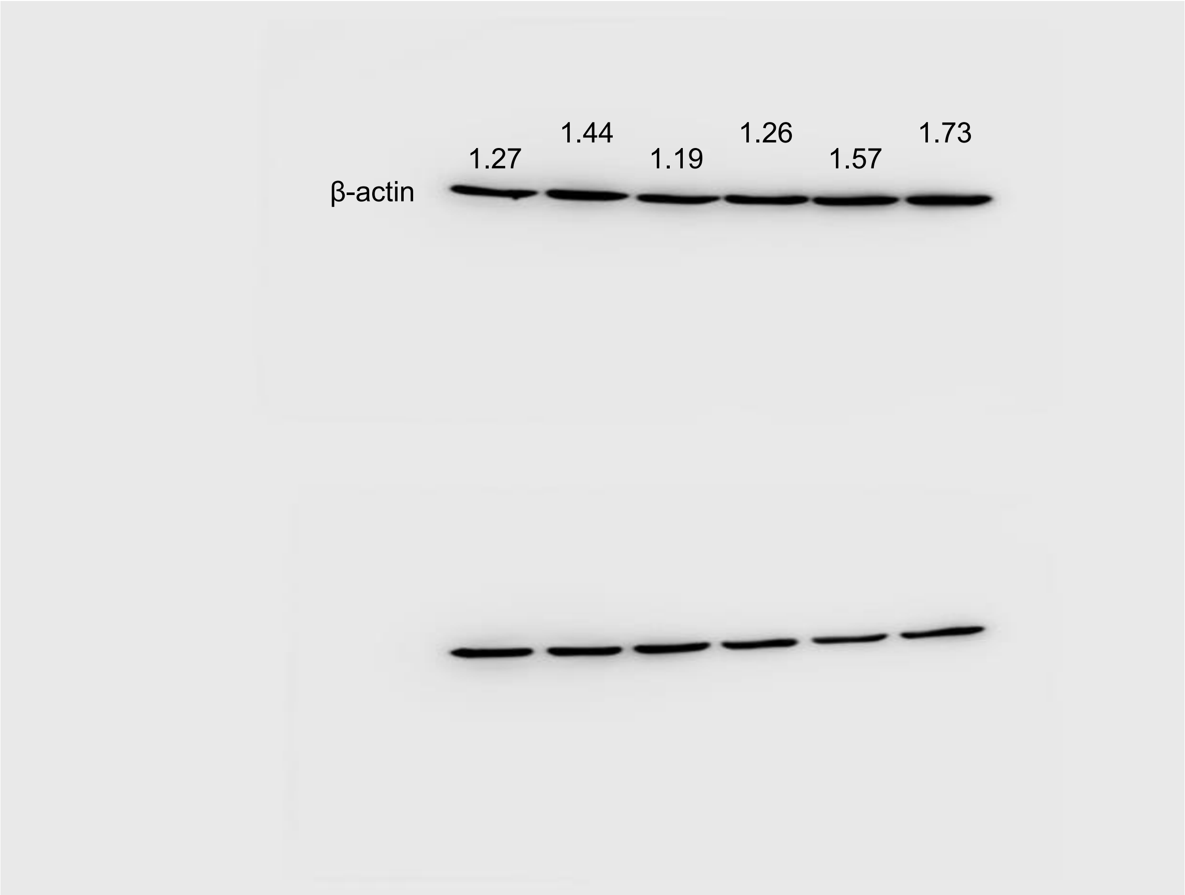
1.27
1.44
1.19
1.26
1.57
1.73
β-actin

### Figure S3

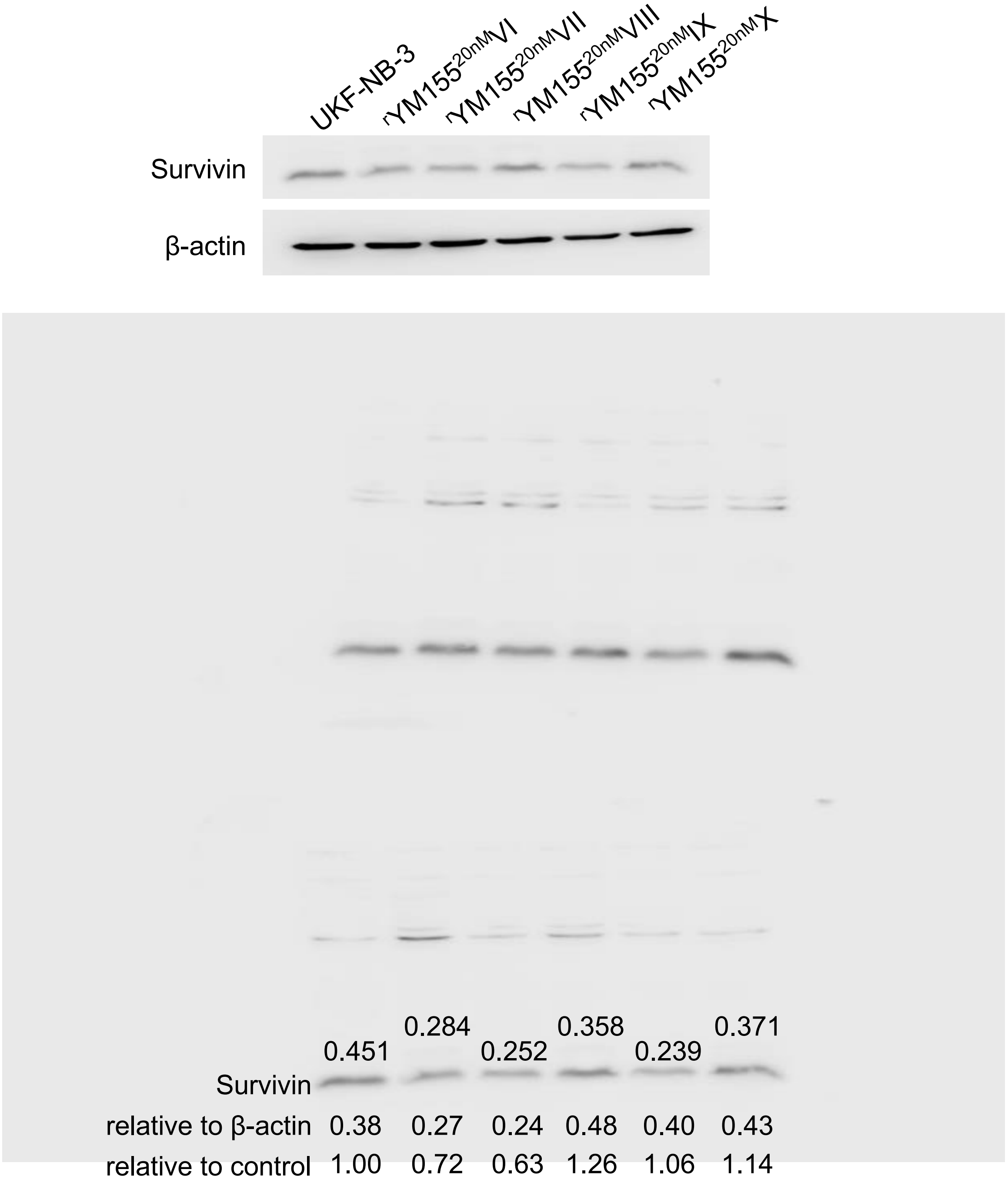
UKF-NB-3
$^{r}$YM155$^{20nM}$VI
$^{r}$YM155$^{20nM}$VII
$^{r}$YM155$^{20nM}$VIII
$^{r}$YM155$^{20nM}$IX
$^{r}$YM155$^{20nM}$X
Survivin
β-actin
0.451 0.284 0.252 0.358 0.239 0.371
Survivin
relative to β-actin 0.38 0.27 0.24 0.48 0.40 0.43
relative to control 1.00 0.72 0.63 1.26 1.06 1.14

### Figure S3

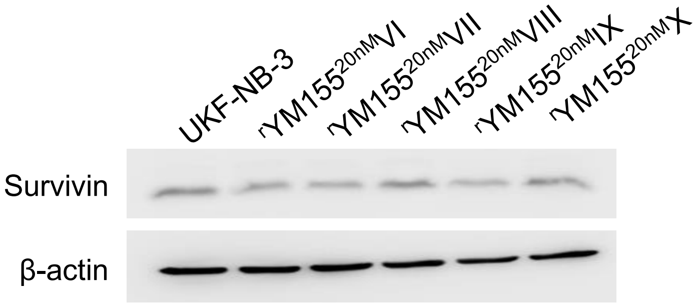

UKF-NB-3
rYM155 20nM VI
rYM155 20nM VII
rYM155 20nM VIII
rYM155 20nM IX
rYM155 20nM X
Survivin
β-actin


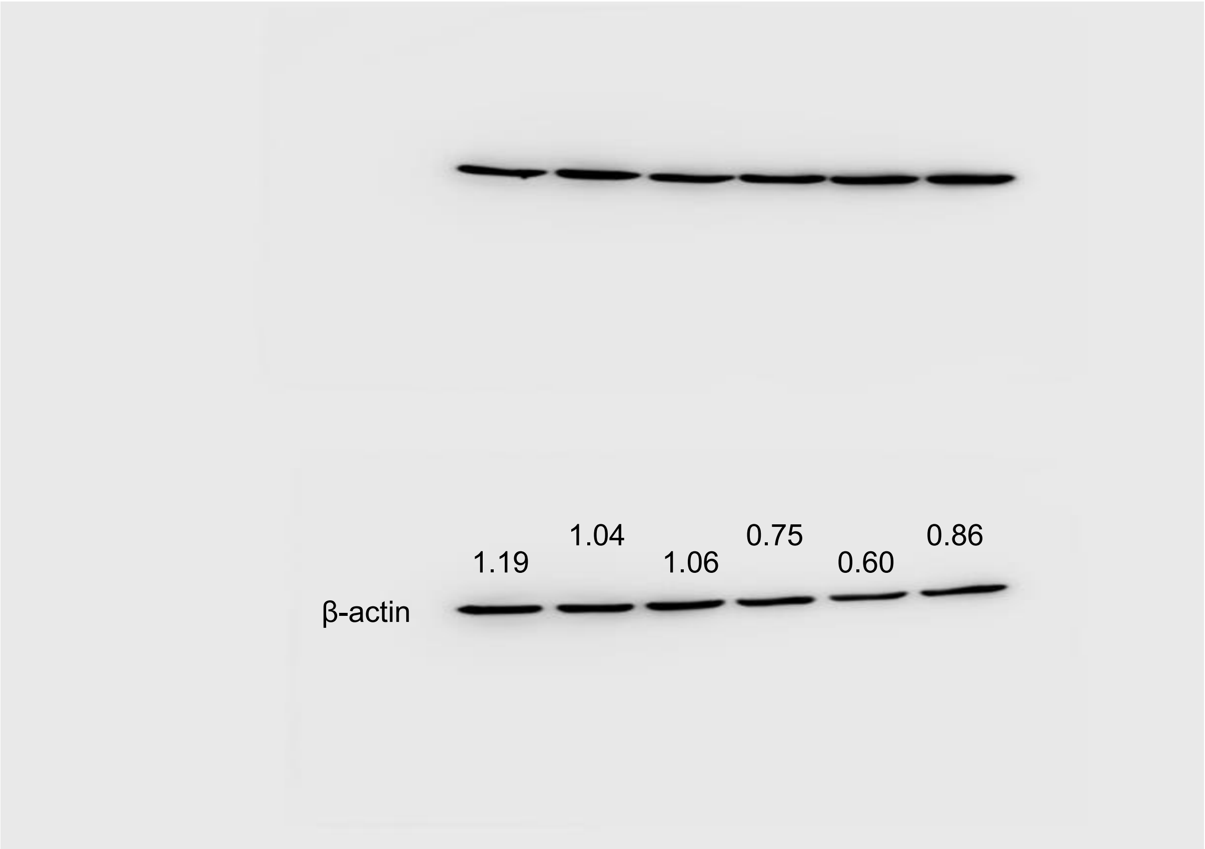

1.19
1.04
1.06
0.75
0.60
0.86
β-actin

**Figure S4**
**survivin**

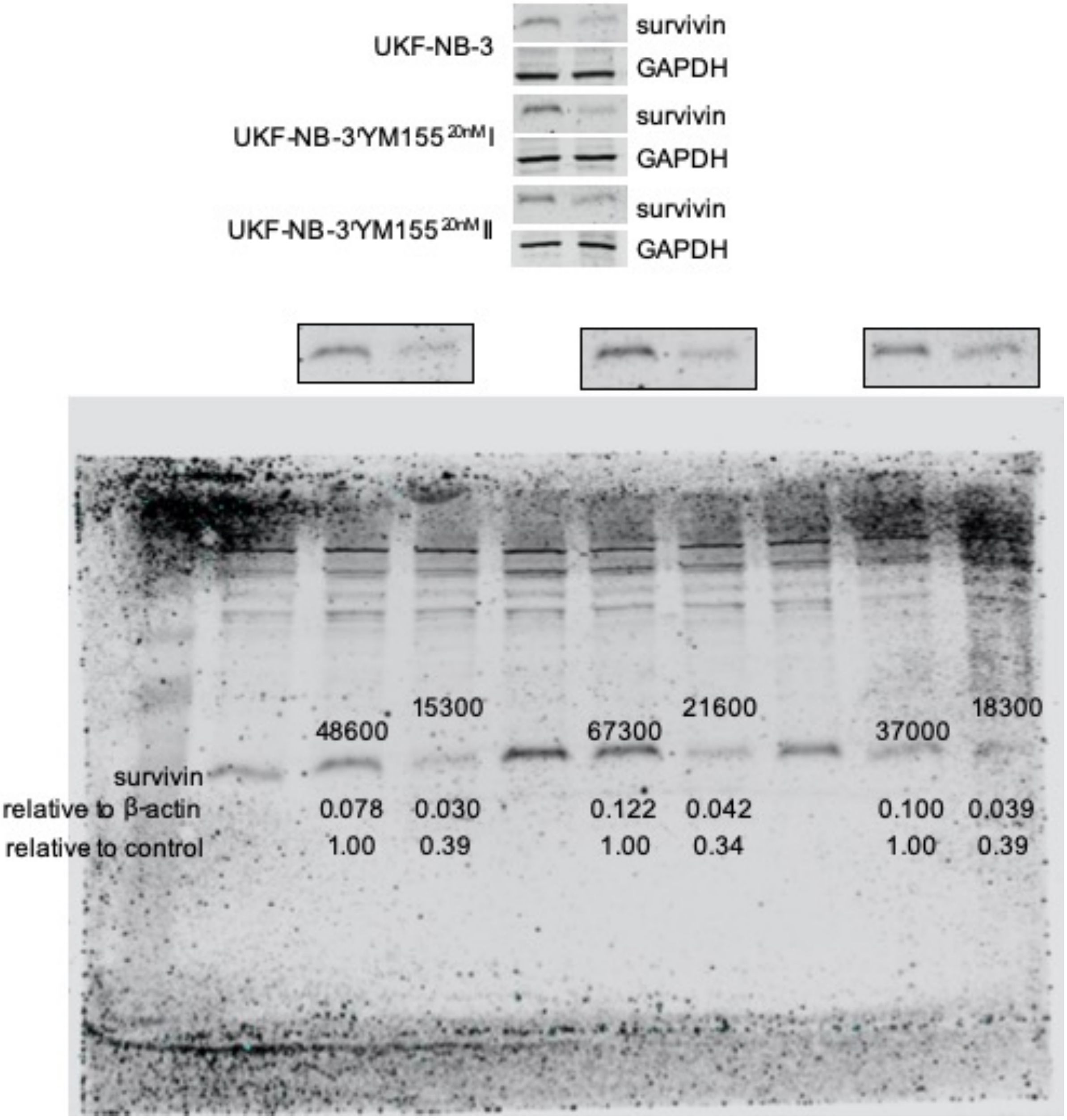

### Figure S4
### GAPDH

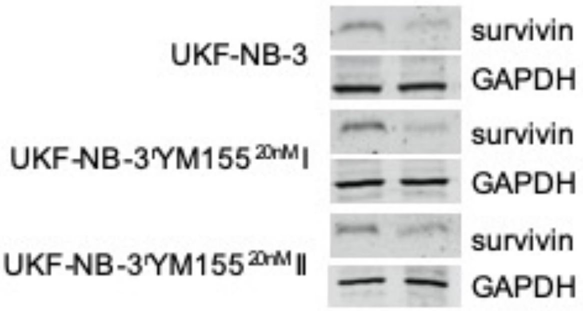


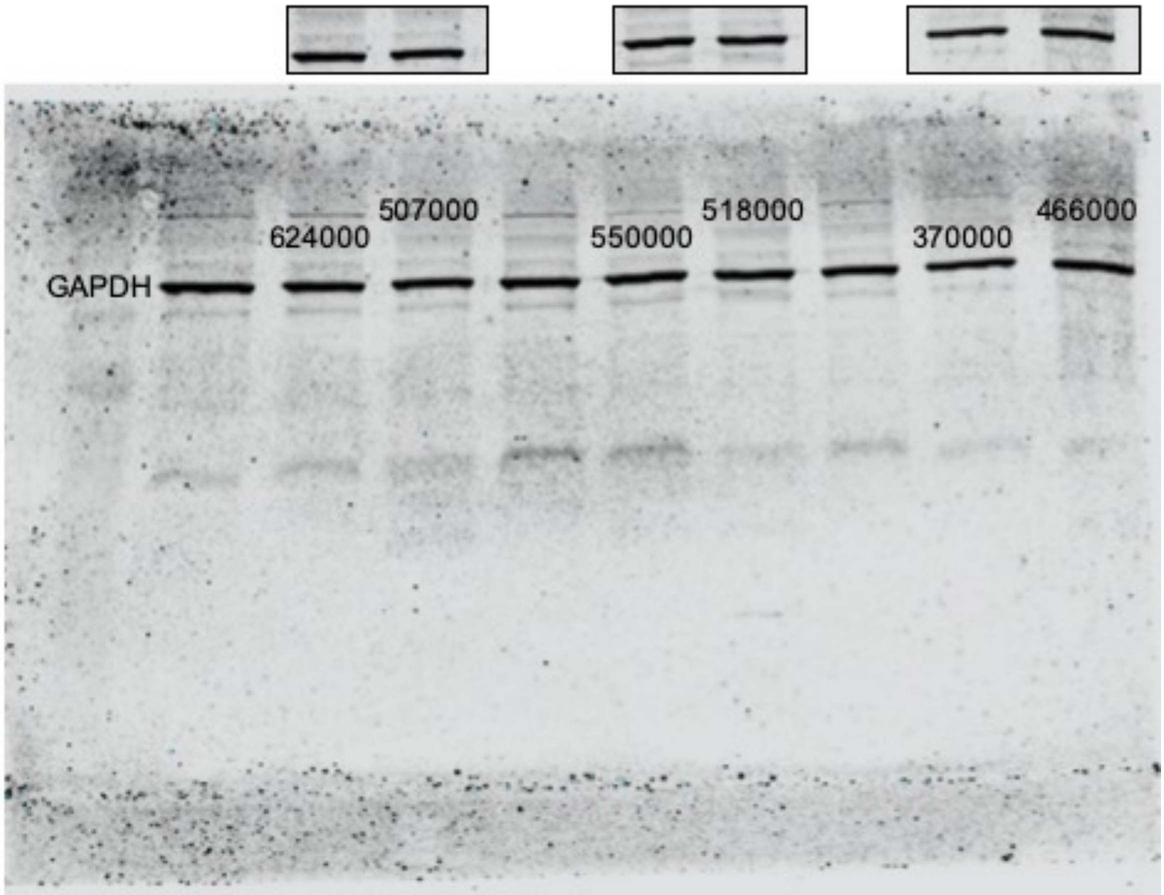

**Figure S4**

**survivin**

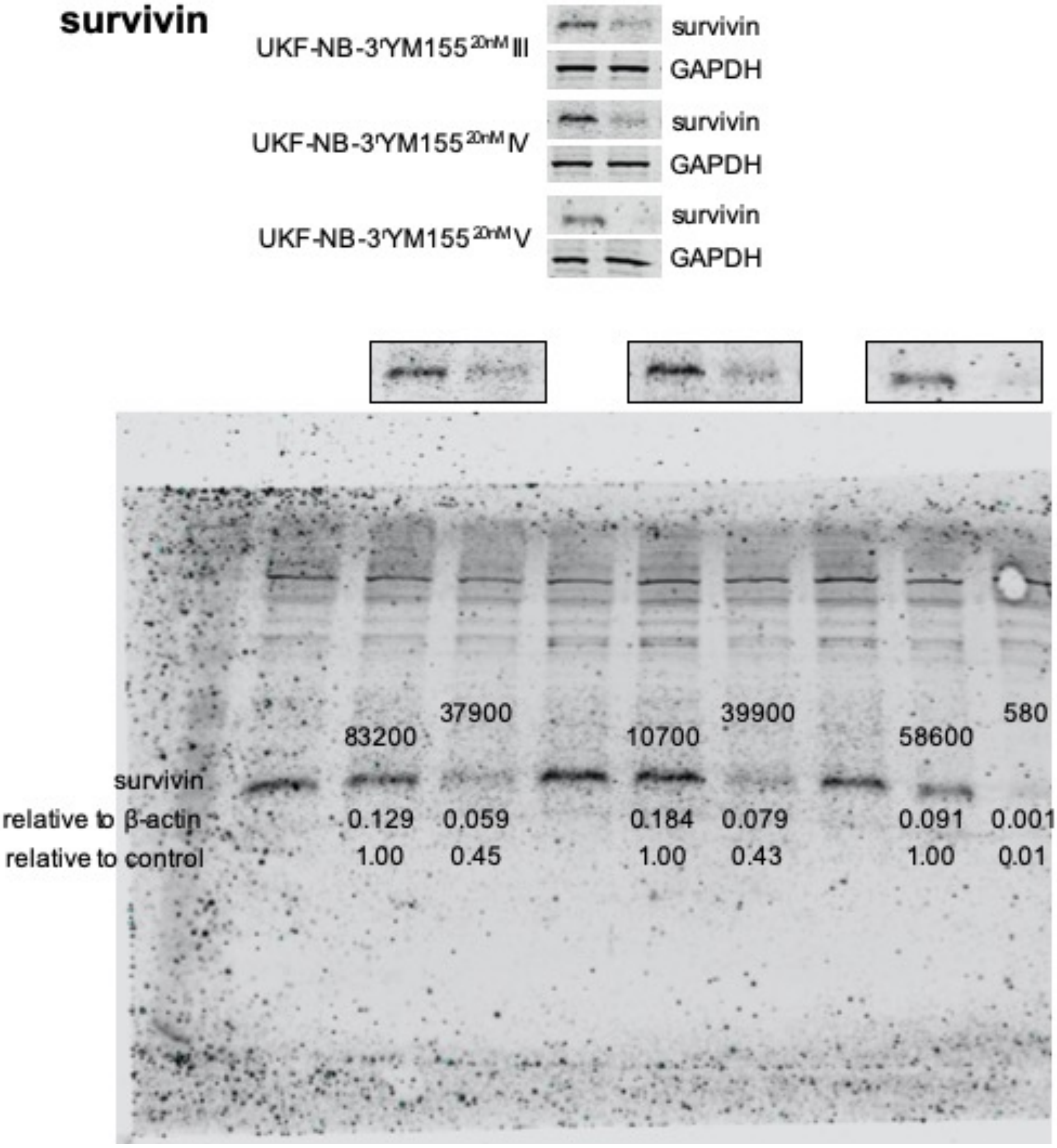

**Figure S4**
**GAPDH**

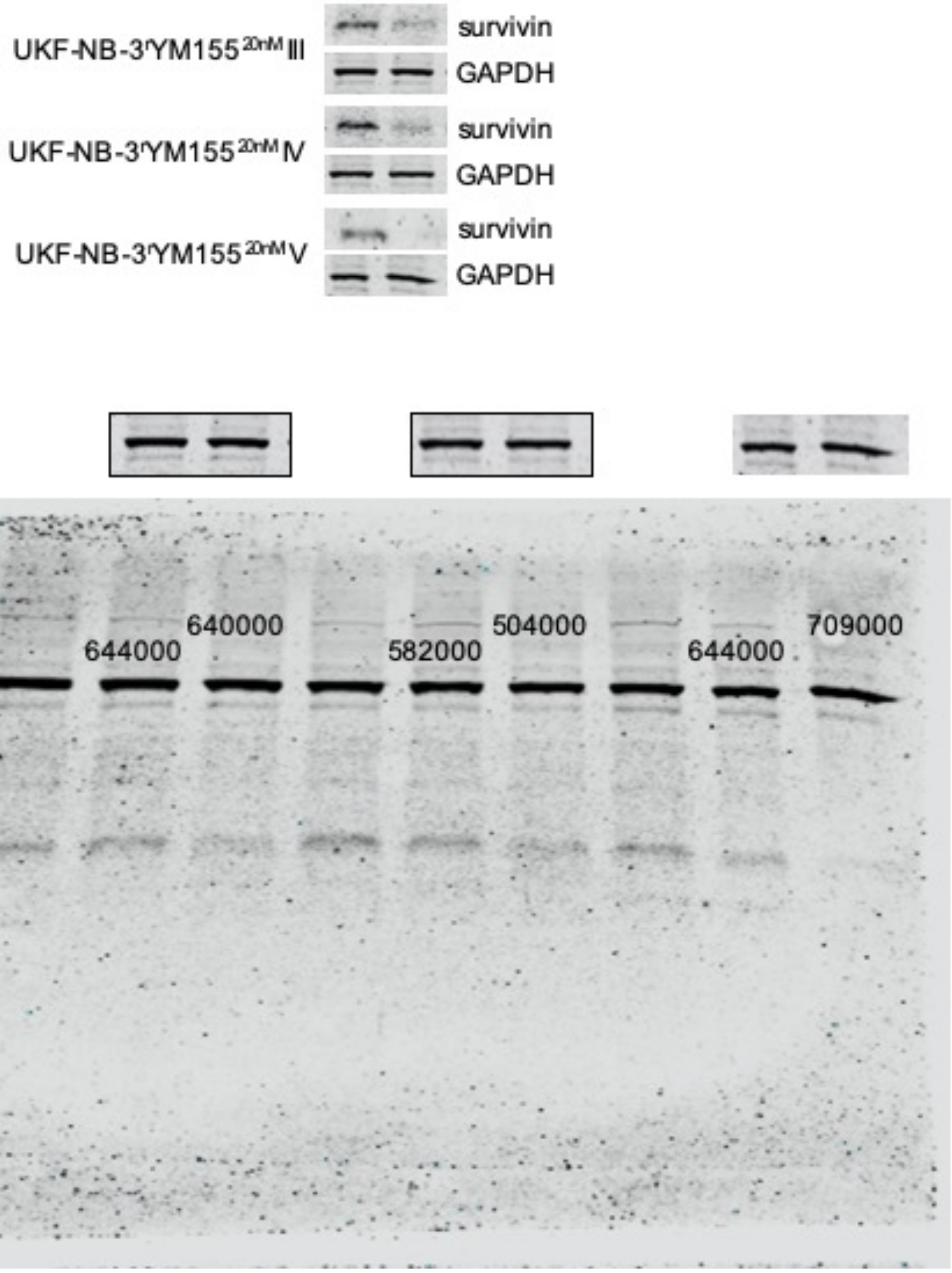

**Figure S4**
**survivin**

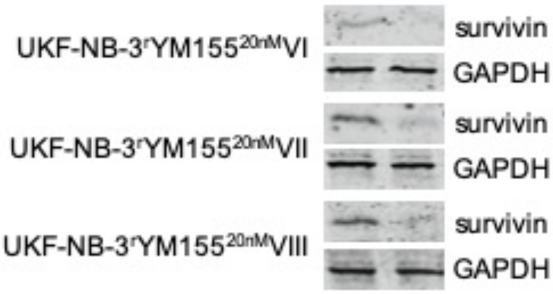


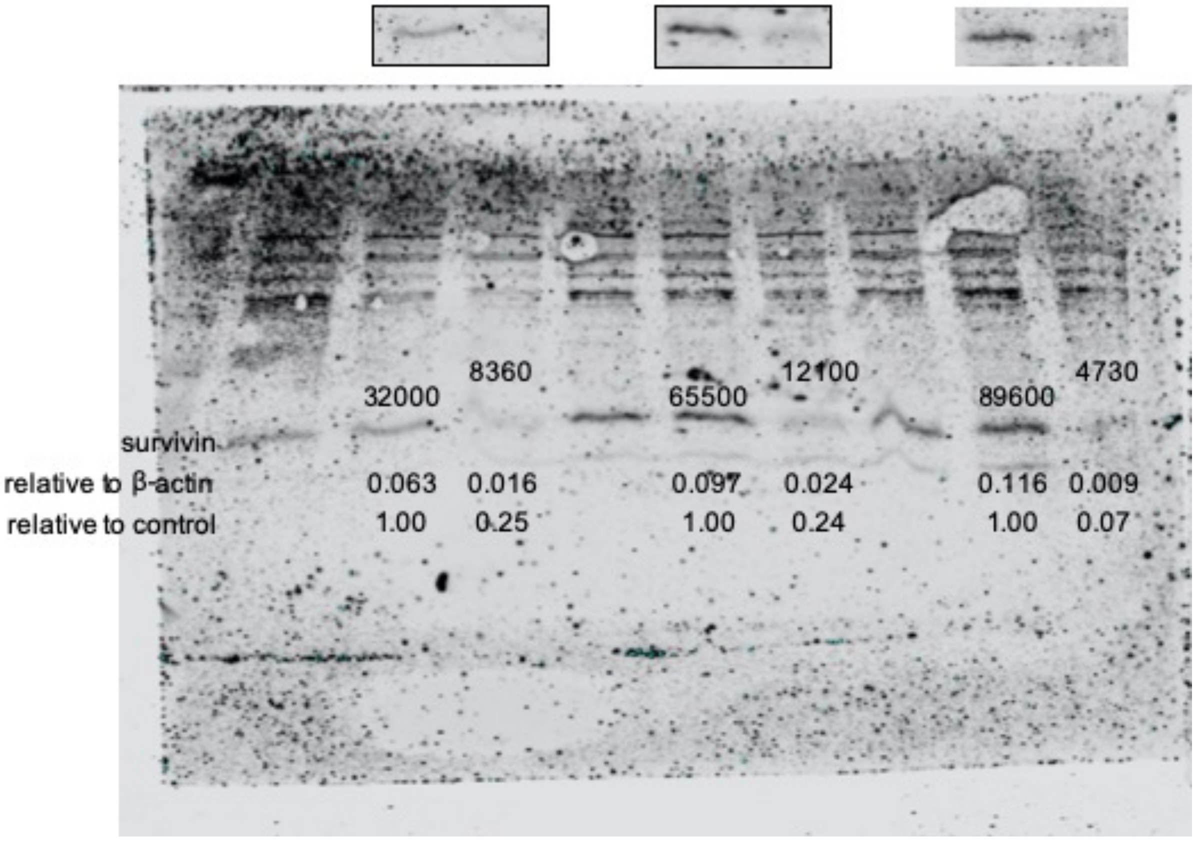

**Figure S4**
**GAPDH**

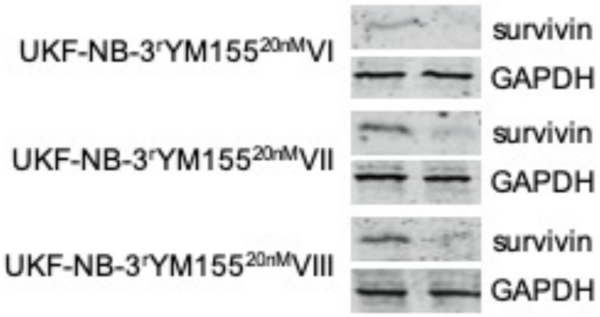


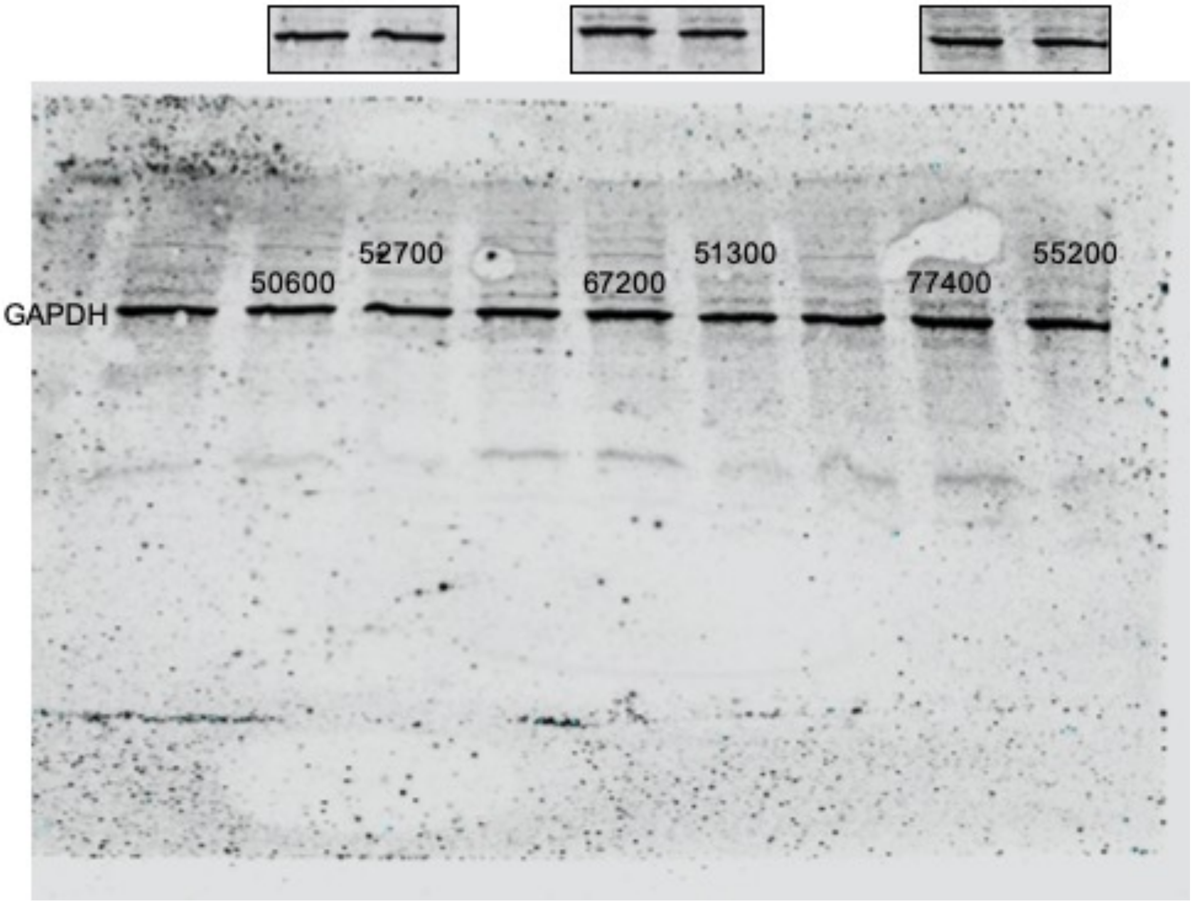

### Figure S4
#### survivin

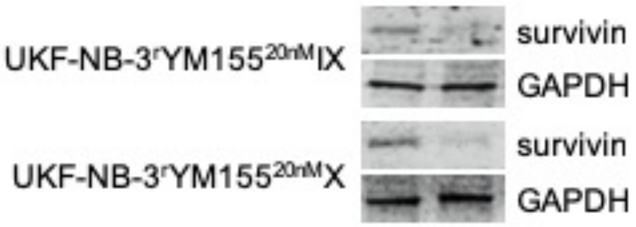


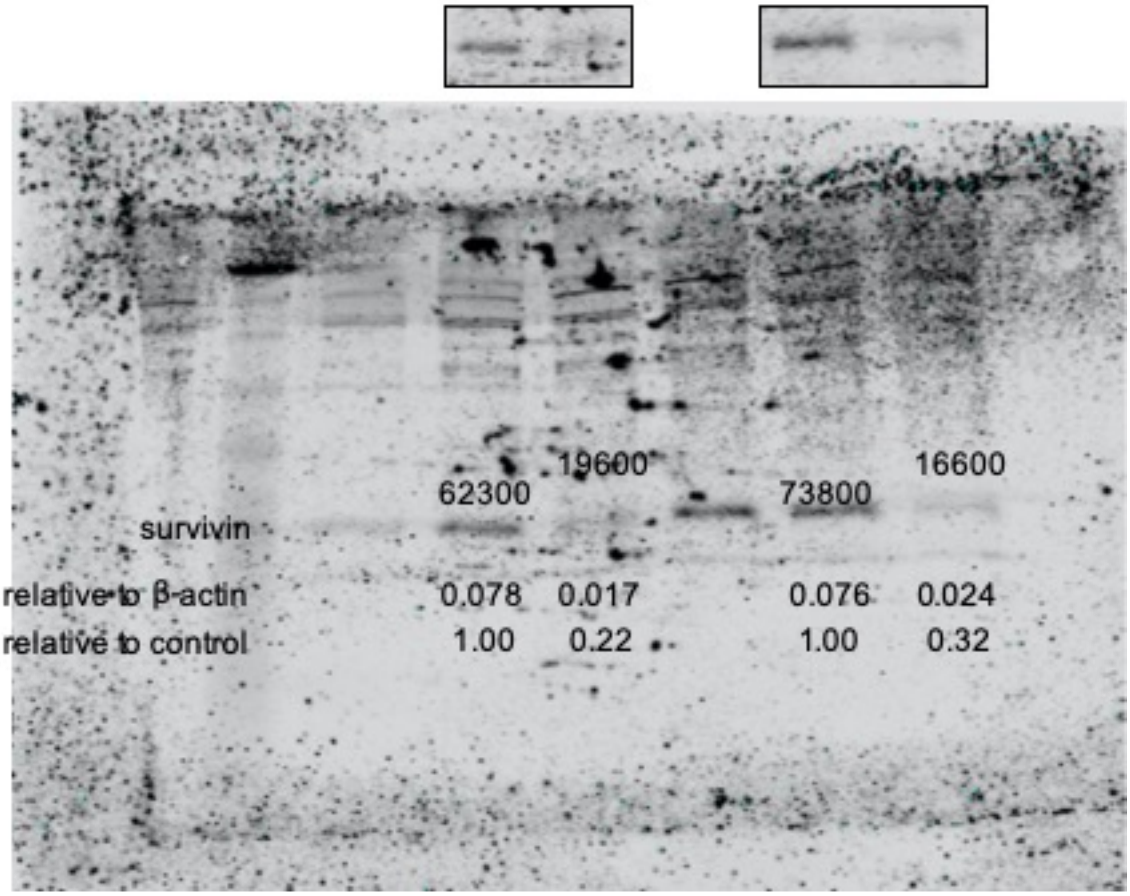

**Figure S4**
**GAPDH**

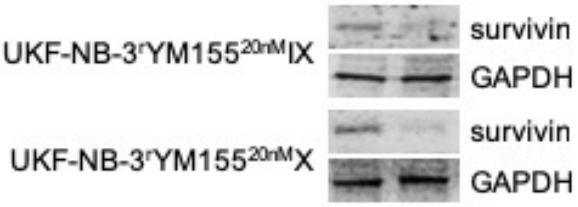


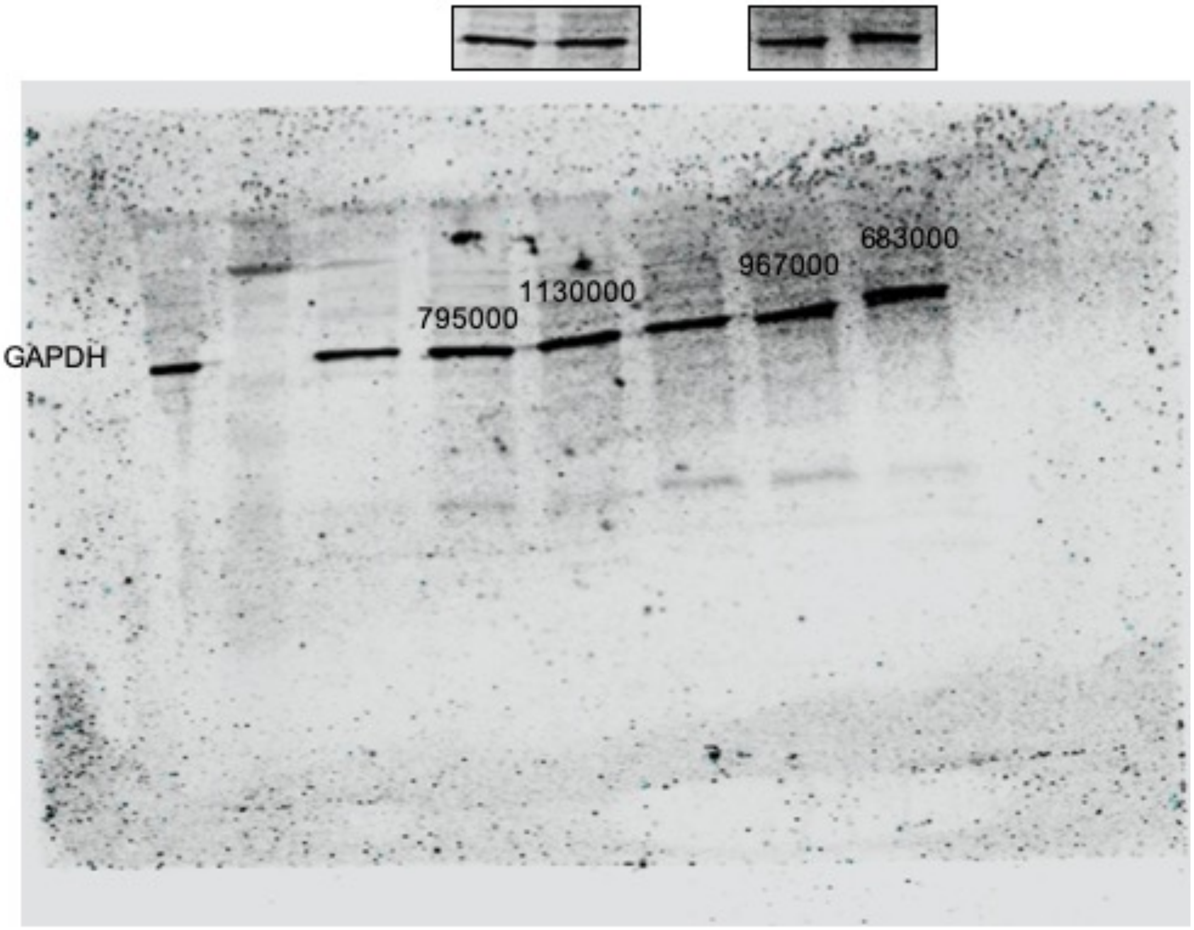


**Figure S4**. Representative Western blots indicating cellular levels of survivin in UKF-NB-3 and its YM155-adapted UKF-NB-3 sub-lines 24 h after transfection with non-targeting siRNA or siRNA directed against BIRC5/ survivin. Densitometric analysis was performed with QuantiOne (BioRad). Survivin levels were normalised to β-actin expression and values relative to control cells are displayed.

**Figure S5**
**ABCB1**

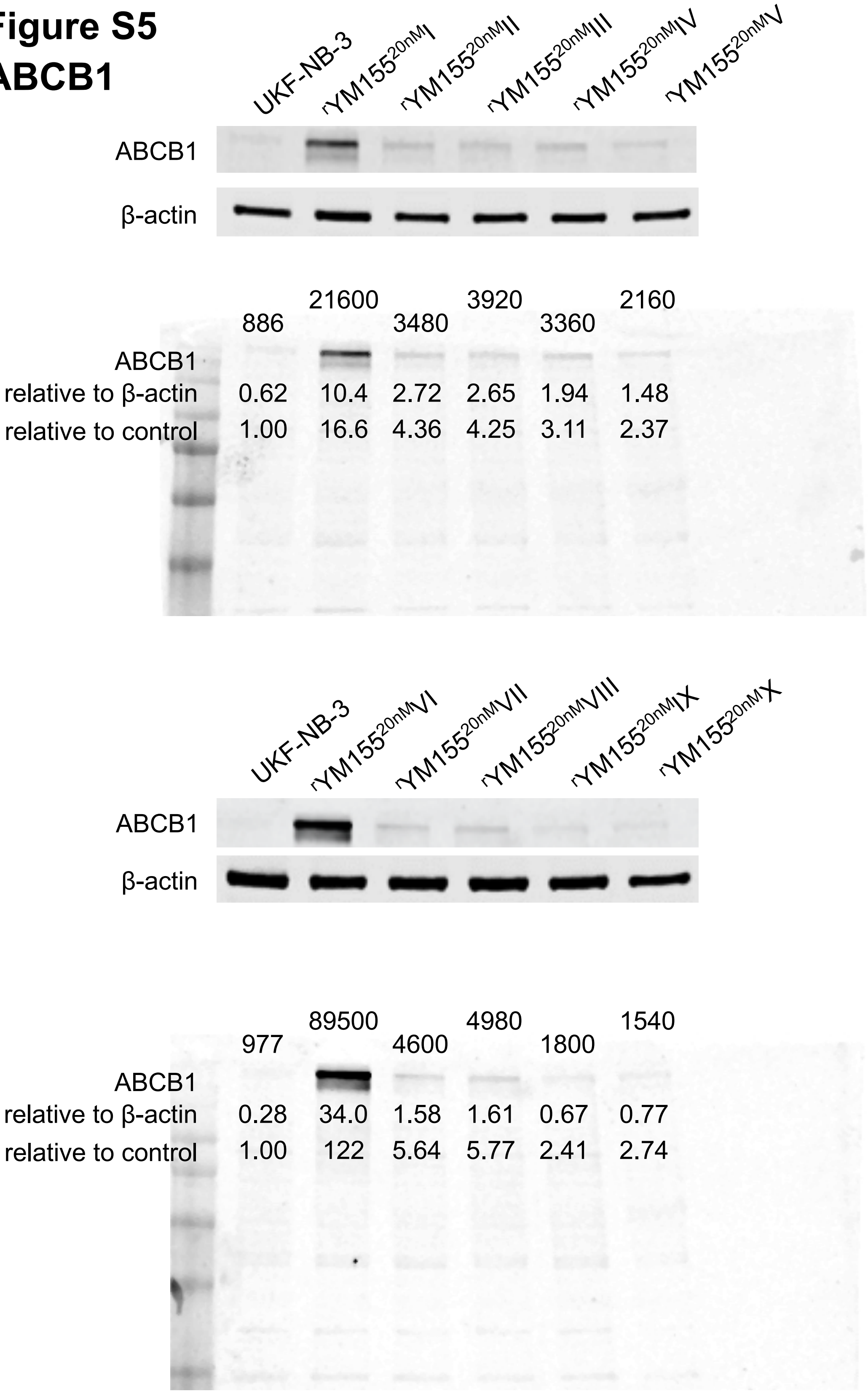

### Figure S5

### SLC35F2

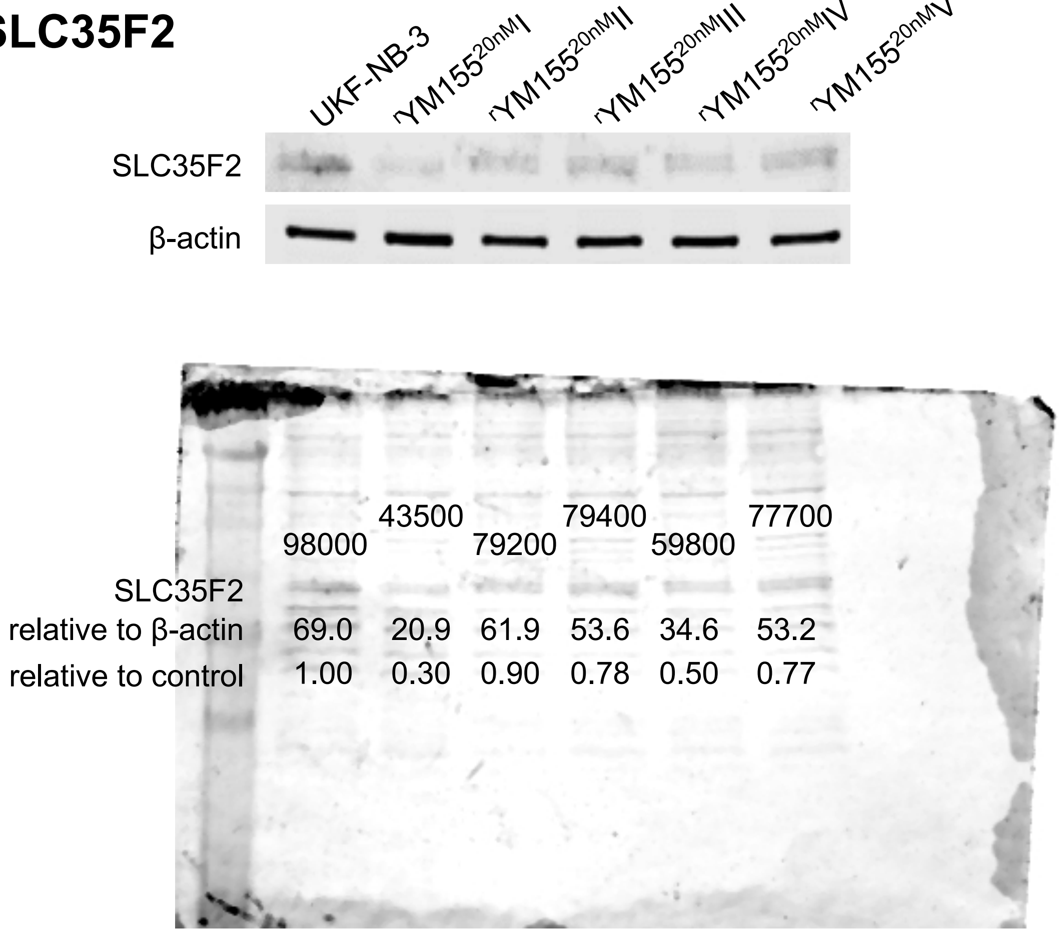

**Figure S5**

**SLC35F2**

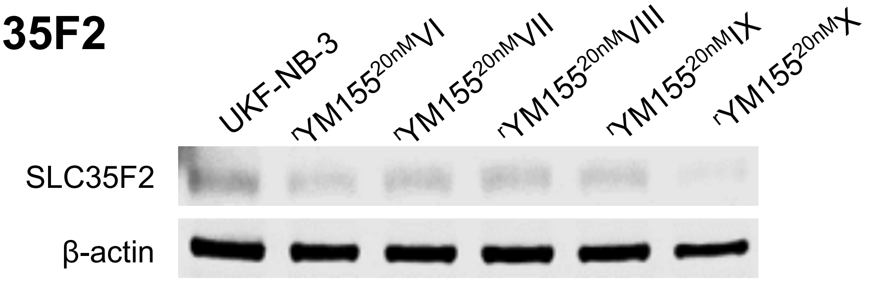


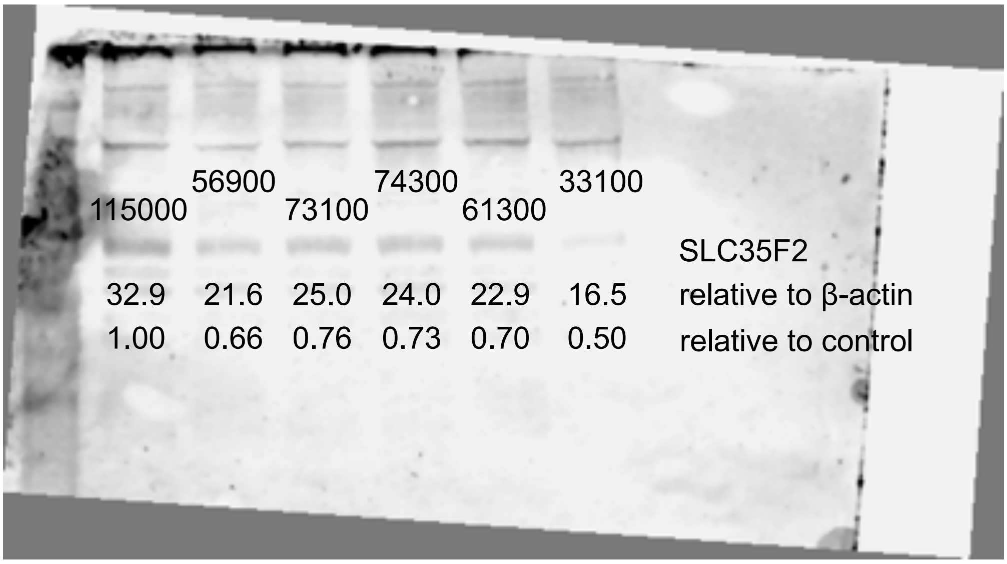

**Figure S5**

**β-actin**

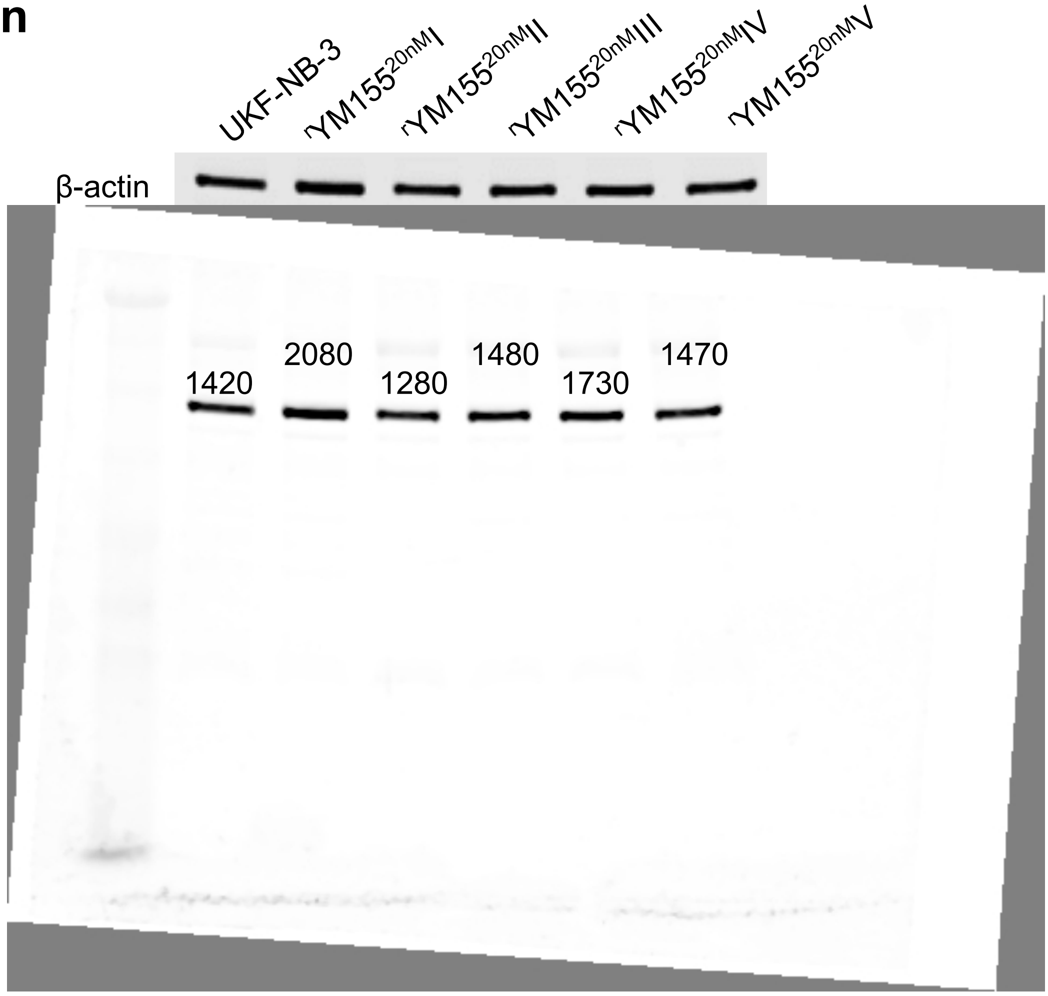

**Figure S5**

**β-actin**

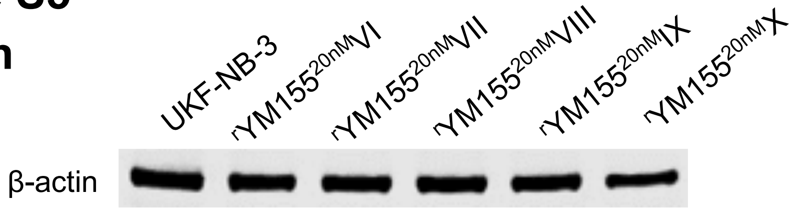


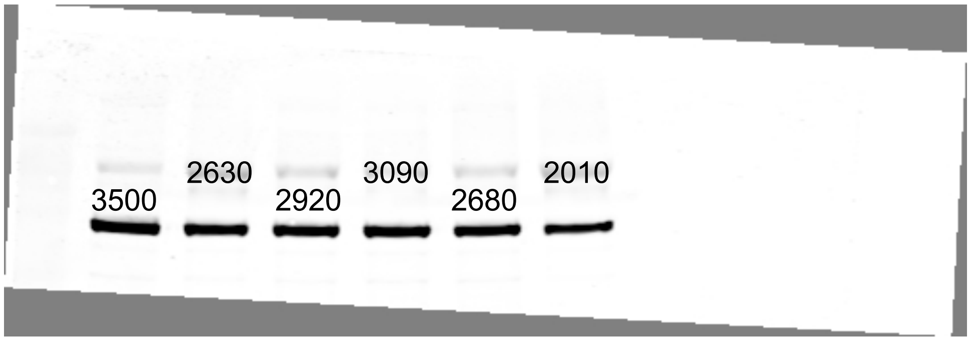


**Figure S5**. Representative Western blots indicating cellular levels of ABCB1 and SLC35F2 in UKF-NB-3 and YM155-adapted UKF-NB-3 sub-lines. Densitometric analysis was performed with QuantiOne (BioRad). ABCB1 and SLC35F2 levels were normalised to β-actin expression and values relative to control cells are displayed.
